# On optimal temozolomide scheduling for slowly growing gliomas

**DOI:** 10.1101/2022.03.10.482967

**Authors:** Berta Segura-Collar, Juan Jiménez-Sánchez, Ricardo Gargini, Miodrag Dragoj, Juan M. Sepúlveda, Milica Pešić, Pilar Sánchez-Gómez, Víctor M. Pérez-García

## Abstract

**Background:** Temozolomide (TMZ) is an oral alkylating agent active against gliomas with a favorable toxicity profile. It is part of the standard of care in the management of glioblastoma, and is commonly used in low-grade gliomas. *In-silico* mathematical models can potentially be used to personalize treatments and to accelerate the discovery of optimal drug delivery schemes.

**Methods:** Agent-based mathematical models fed with either mouse or patient data were developed for the *in-silico* studies. The experimental test beds used to confirm the results were: mouse glioma models obtained by retroviral expression of EGFR wt or EGFR vIII in primary progenitors from p16/p19 ko mice and grown *in vitro* and *in vivo* in orthotopic allografts, and human glioblastoma U251 cells immobilized in alginate microfibers. The patient data used to parametrize the model were obtained from the TCGA/TCIA databases and the TOG clinical study.

**Results:** Slow growth ‘virtual’ murine gliomas benefited from increasing TMZ dose separation *in silico*. In line with the simulation results, improved survival, reduced toxicity, lower expression of resistance factors and reduction of the tumor mesenchymal component were observed in experimental models subject to long-cycle treatment, particularly in slowly-growing tumors. Tissue analysis after long-cycle TMZ treatments revealed epigenetically-driven changes in tumor phenotype, which could explain the reduction in glioma growth speed. *In-silico* trials provided support for methods of implementation in human patients.

**Conclusions:** *In-silico* simulations, and *in-vitro* and *in-vivo* studies show that TMZ administration schedules with increased time between doses may reduce toxicity, delay the appearance of resistances and lead to survival benefits mediated by changes in the tumor phenotype in gliomas.

**IMPORTANCE OF THE STUDY:** *In-vivo* evidence is provided of improvements in survival, resistance, and toxicity from TMZ schemes with long rest periods between doses in slowly-growing GBM mouse models. The results match hypotheses generated *in silico* using a mathematical model incorporating the main biological features and fed with real patient data. An epigenetically-driven change in tumor phenotype was also revealed experimentally, which could explain the reduction in glioma growth speed under the ‘long cycle’ scheme. To determine the extent to which our results hold for human patients, large sets of simulations were performed on virtual patients. These *in-silico* trials suggest different ways to bring the benefits observed in experimental models into clinical practice.

## INTRODUCTION

Adult gliomas are the most common primary malignant tumors of the central nervous system. A novel classification of these neoplasms has been proposed^1–3^, using a combination of molecular and histopathological features,. Especially important for patient prognosis is the presence of mutations in the isocitrate dehydrogenase 1/2 (IDH1/2) genes, which distinguishes IDH mutant (IDHmut) gliomas from IDH wild-type (IDHwt) glioblastomas (GBM)^4^. GBMs are diagnosed at a later age (median 64) and are characterized by the presence of additional alterations. Some of them are mutations in the telomerase reverse transcriptase (TERT) promoter, amplification of the epidermal growth factor receptor (EGFR), and gains and losses of chromosomes 7 and 10, respectively^5^. GBMs have a dismal prognosis (15 months’ overall survival) despite the standard-of-care treatment, which consists of maximal surgical resection followed by radiotherapy (RT) plus concomitant and adjuvant chemotherapy (CT) with temozolomide (TMZ), an oral alkylating agent^6,7^. TMZ is administered orally at a dose of 75 mg/m^2^ daily throughout RT, plus six cycles of maintenance TMZ 150–200 mg/m^2^ for 5 out of 28 days^6^. The cytotoxicity of this drug is attributed to the addition of methyl groups to DNA, and especially to the formation of O6-methylguanine (O6-meG) lesions and the subsequent formation of double-strand breaks during DNA replication, which requires cell division for the emergence of the cytotoxicity. As O6-meG can be removed by methylguanine methyltransferase (MGMT) in tumors expressing this protein, MGMT promoter methylation is considered a predictive biomarker of TMZ response in gliomas^8^. However, challenges remain in establishing reliable inter-laboratory assays^9^, as well as in estimating the effect of limited MGMT promoter methylation on outcomes^10^. Moreover, based on the short half-life of TMZ, it was suggested that high doses or repeated doses could improve the effect of this commonly-used drug, and reduce the capacity of cells to repair the DNA^11^. However, dose-dense TMZ, with the administration of lower but continuous doses with the aim of depleting intracellular MGMT, did not show improved efficacy in newly diagnosed GBM,^12^ and has shown only modest results in recurrent tumors^13–16^ at the cost of increased hematological toxicity. Simple mathematical models of lower grade gliomas (LGG) (grade 2/3 gliomas) have suggested that TMZ schemes with longer rest periods between cycles could improve the survival of patients^17^. However, no study has tested the efficacy of longer spacing between cycles/doses. Therefore, there may be room for improvements in the schedules used for TMZ administration to glioma patients.

Mathematical models describe real systems by abstraction and mathematical formalism. They enable extrapolation beyond the situations originally analyzed, allowing for quantitative predictions, inference of mechanisms, falsification of underlying biological hypotheses, and quantitative descriptions of relationships between different components of a system. They cannot replace experimental results obtained by biomedical models, but may complement experimentation in providing a broader picture^18^. Moreover, in the field of oncology they can suggest what the best RT, CT or combination regimens might be, aiding the implementation of the treatment of different cancers, including gliomas. A number of mathematical models of LGGs have been constructed in order to study the optimal delivery scheme of cytotoxic therapies^17,19,20,21^. It is important to emphasize that in slow-growth gliomas, such as LGGs, only a small percentage of cells is proliferating as shown by fractions of Ki67 positive cells typically below 5%^22^. Notably, even in GBM, there is a small group of tumors with reduced Ki67%^23^. It may be guessed that intensive therapies intended to deliver the maximum tolerated dose in the shortest possible time may be overkill for these tumors. Indeed, several authors have proposed that schemes with longer spacing between doses could produce better results in LGG patients^17,20,21,24^. However, the simple mathematical models developed previously account only for a limited number of key biological processes. Notably, no detailed theoretical models, including realistic resistance mechanisms, have been considered previously. Also, it is still unknown whether the potential gain observed *in silico* for LGGs would apply to GBMs as well. First of all, the proneural (PN) to mesenchymal (MES) phenotypic transition observed in GBMs, either spontaneously or as a result of (CT/RT), would limit the effect of TMZ as the tumor becomes more resistant^25,26^. Also, from the biological point of view, there is a growing body of literature suggesting that the evolution of tumor cells to a fully drug-resistant state may often proceeds through a reversible drug-tolerant phase^27^, so-called persister cells^28^. Rabé et al identified a population with persister characteristics under TMZ treatment of glioma cells^29^. One may guess that longer spacing between cycles could delay the PN-MES transition, as well as allowing for persister cells to revert to their normal sensitive states. Neither of these facts has been accounted for by previous mathematical modeling approaches.

The intention of this paper was to set out a proof of concept supporting treatment regimes with longer times between doses of TMZ (hereafter referred to as protracted temozolomide schedules, PTS). Our hypothesis was that PTS could lead to reduced appearance of persister cells. Moreover, reduced toxicity and increased tolerance was anticipated from increasing the rest period between cycles. To test these ideas, a stochastic mesoscopic discrete mathematical model was first produced, including all relevant biological processes expected to play a role in the response of GBMs to TMZ. Next, PTS was studied *in vitro* and in animal models and found to be in good agreement with the *in silico* observations. The tissue of TMZ-responding allografts was then analyzed, and showing a striking change in tumor phenotype, with an increase in PN and a decrease in MES markers, driven by epigenetic changes, which could explain the reduction in tumor growth. Finally, exploratory virtual ‘clinical’ trials were performed with *in-silico* tumors simulated with the previous mathematical model, to guide the implementation of the concept in clinical practice. The results indicated that survival is increased as spacing between doses becomes progressively larger, until a threshold is reached; spacings beyond this threshold would fail to improve survival.

## METHODS

### Cell lines and cell culture

The human GBM U251 cell line was purchased from American Type Culture Collection (ATCC, USA). It was cultured in Dulbecco’s Modified Eagle Medium (Biological Industries, USA) supplemented with 10% fetal bovine serum (Sigma-Aldrich, Germany), 2 mM glutamine (Sigma-Aldrich, Germany), 5,000 U/ml penicillin, and 5 mg/ml streptomycin (GibcoTM, Thermo Fisher Scientific, United States). Cells were cultivated at 37°C in a humidified 5% CO_2_ atmosphere and passaged twice a week after reaching 80-90% confluence using 0.25% trypsin/EDTA.

Mouse SVZ cell lines were obtained by retroviral expression of EGFRwt or EGFRvIII (pBabe-EGFR wt (#11011) and MSCV-XZ066-GFP-EGFR vIII (#20737)) in primary neural stem cell cultures obtained from the subventricular zone (SVZ) of p16/p19 ko mice^30^. After infection, the cells were injected into nude mice, and the tumors that grew were dissociated and the lines SVZ-EGFRwt/amp and SVZ-EGFRvIII were established^31^. Both models express GFP and luciferase as a reporter. Cells were grown as previously described^30,31^. Briefly, they were maintained in stem cell medium; Neurobasal (Invitrogen) supplemented with B27 (1:50) (Invitrogen); GlutaMAX (1:100) (Invitrogen); penicillin-streptomycin (1:100) (Lonza); 0.4% heparin (Sigma-Aldrich); and 40 ng/ml EGF and 20 ng/ml bFGF2 (Peprotech). For dissociation and passage Accumax (ThermoFisher) was used.

### Production of alginate microfibers with U251 immobilized cells

Alginate microfibers with cells were produced by extrusion as described earlier^32^. U251 cell lines were immobilized in alginate microfibers by the same procedure. Briefly, 4 × 10^6^ cells/ml were mixed with a 2 % w/v Na-alginate solution to obtain final concentrations of 1.5% w/v Na-alginate. The Na-alginate solution with cells was manually extruded through a blunt edge stainless steel 25G needle immersed in the gelling bath (3 % w/v Ca(NO_3_)2 x 4H_2_O). Due to the exchange of Na^+^ with Ca^2+^, the liquid stream solidified in the gelling bath, thus forming insoluble microfibers. The microfibers were left in the bath for 15 min in order to complete gelling and were then washed with medium. After cell immobilization, 0.5 g of alginate fibers were distributed into a T25 flask and cultured for 28 days without passage in 13 ml of MEM medium. 50% of the medium was changed twice a week.

### Viability study

The impact of three different TMZ (Sigma-Aldrich) (100 μM) treatment modalities (everyday treatments, X+1, and protracted (every 3 days, X+3; every 7 days, X+7)), starting from day 7, was determined by comparing the effects on cell viability, morphology, and aggregation using a CalceinAM (CAM)/propidium-iodide (PI) assay. U251 cells immobilized in alginate microfibers were cultured for 28 days and stained using CAM/PI as a LIVE/DEAD staining. Alginate microfibers containing cells were incubated for 45 min at 37 °C in medium with CAM in a final concentration of 4 μM while PI was added to a final concentration of 5 μM. Fluorescence microscopy images were taken using a Leica TCS SP5 II Basic confocal laser-scanning microscope (Leica Microsystems CMS GmbH; Germany), visualizing live (green) and dead (red) cells at every z-axis encompassing the alginate microfiber. Z-stack projections and quantitative estimation of the cell mass were analyzed using ImageJ software.

### *In-vitro* treatments of mouse cells

SVZ EGFR wt/amp cells were incubated in the presence of TMZ (25 μM), which was supplemented three times: one day (X+1), three days (X+3) or seven days (x+7) after the first dose. In a different experiment, SVZ EGFR wt/amp cells were treated with TMZ (25 μM) and/or Azacytidine (AZA) (Sigma-Aldrich) (5 μm) for 8 days. For both experiments, cells were collected and lysed and subsequently analyzed by qRT-PCR, as described below.

### Intracranial tumor formation and treatment *in vivo*

Animal experiments were reviewed and approved by the Research Ethics and Animal Welfare Committee at “Instituto de Salud Carlos III” (PROEX 02/16), in agreement with the European Union and national directives. Intracranial transplantation to establish orthotopic allografts was performed injecting 300,000 cells (resuspended in 2 μl of culture cell medium) with a Hamilton syringe into athymic Nude-Foxn1nu brains (Harlan Iberica). Female mice (2-3 months of age) were used, 7 to 10 animals per group. The injections were made into the striatum (coordinates: A–P, −0.5 mm; M–L, +2 mm, D–V, −3 mm; related to Bregma) using a Stoelting Stereotaxic device. Mice were treated with TMZ (10 or 50 mg/kg through intraperitoneal injection, i.p., with the schedules given in the various experiments). TMZ was dissolved in PBS+1% BSA, which was used to treat control animals.

### Immunohistochemical (IHC) staining

Tumor samples were fixed in 10% formalin overnight, dehydrated through a series of graded ethanol baths and then infiltrated with paraffin. Then, 2.5 μm-thick sections were obtained in a microtome and then sections were rehydrated and permeabilized (1% triton X-100). Antigen retrieval was performed with Citrate Buffer (10 mM, pH 6) in a pressure cooker (2 min). Endogenous peroxidase inhibition and blocking with normal horse serum was also carried out before the incubation with primary antibodies (anti-rabbit caspase3, 1:100, Cell signaling #9662), anti-mouse ki67, 1:100 Dako #M7248) (overnight, 4 °C) and biotinylated secondary antibodies (HRP anti-mouse and HRP anti-rabbit, 1:200, GE Healthcare (2h at room temperature). Target proteins were detected with the ABC Kit and the DAB kit (Vector Laboratories).

### Western Blot analysis

For protein expression analysis, mouse tumor tissue was processed by mechanical disruption in a lysis buffer (Tris–HCl pH 7.6, 1mMEDTA, 1mMEGTA, 1% SDS, and 1% Triton X-100) followed by heating for 15 min at 100°C. Protein content was quantified by using a BCA Protein Assay Kit (Thermo Fisher Scientific). Approximately 30 μg of proteins were resolved by 10% or 12% SDS-PAGE, and these were then transferred to a nitrocellulose membrane (Hybond-ECL, Amersham Biosciences, Little Chalfont, UK). The membranes were blocked for 1 h at room temperature in TBS-T (10 mM Tris–HCl (pH 7.5), 100 mM NaCl, and 0.1% Tween-20) with 5% skimmed milk, and then incubated overnight at 4°C, with the corresponding primary antibody (mouse anti-MGMT 1:1000, BD Biosciences, #557045), mouse anti-GAPDH (1:1.500, Santa Cruz Biotechnology #sc-47724), rabbit anti-pTyr1068-EGFR (1:1.000, Cell Signaling #3777) and rabbit anti-phospho-NF-kB p65 (Ser536) (1:1000, Cell Signaling #3033) diluted in TBS-T. After being washed 3 times with TBS-T, the membranes were incubated for 2 h at room temperature with their corresponding secondary antibody (HRP-conjugated anti-mouse (#NA931) or anti-rabbit (#NA934), Amersham Biosciences) diluted in TBS-T.

### RNA extraction and RT-PCR

The impact of three different TMZ treatment modalities was determined by comparing the effects on gene expression in human glioma cells and in the mouse models.

- Human glioma cells: After 28 days of incubation, alginate microfibers with U251 cells were dissolved and cells were released. To release cells, alginate microfibers were dissolved in 0.5 mM EDTA for 10 min at 37°C. Total RNA was extracted from control and treated group of cells. The extractions were carried out using Tri Reagent Solution (Invitrogen LifeTechnologies, USA) according to the manufacturer’s instructions. cDNA was synthesized using 2 μg total RNA and High-capacity cDNA reverse transcription kit (Applied Biosystems, USA) according to the manufacturer’s instructions.
- Mouse samples. Brain tumors were dissected out after the mouse sacrifice and fresh frozen. Alternatively, mouse cells grown *in vitro* were collected and fresh frozen. RNA was extracted from the tissue or the cells using the RNA isolation Kit (Roche). Total RNA (1μg) was reverse transcribed with PrimeScript RT Reagent Kit (Takara).

Quantitative real time PCR was performed using the Light Cycler 1.5 (Roche) with the SYBR Premix Ex Taq (Takara). The primers used for each reaction are indicated in Supplementary Table S1. All experiments were performed in triplicate and relative gene expression levels were analyzed by the 2-ddCt method^33^.

### *In-silico* analysis

The Cancer Genome Atlas (TCGA) GBM dataset was accessed via UCSC xena-browser (https://xenabrowser.net) to extract proliferation gene expression levels. Classification into classical, mesenchymal, neural and proneural subtypes was retrieved from the TCGA GBM data set^5^, together with gene expression values. Differences in gene expression between different groups were calculated using Student’s t-test.

### Statistical analysis

The difference between experimental groups was assessed by paired *t*-test and one-way analysis of variance (ANOVA). For Kaplan-Meier survival curves, the significance was determined by the two-tailed log-rank test. All analyses were carried out with the GraphPad Prism 5 software. *P* values below 0.05 were considered significant (\**P* < 0.05; \*\**P* < 0.01; \*\*\**P* < 0.001; \*\*\*\**P* < 0.0001; n.s., not significant), both for mouse and simulated tumors. All experimental quantitative data presented are the means +/− SEM from at least three samples or experiments per data point.

### Discrete mathematical model

An on-lattice agent-based mesoscopic model^35^ was adapted to simulate the longitudinal growth dynamics of glioma and its response to treatments *in silico.* A comprehensive model description is provided in the Supplementary Information. This study included three basic cellular populations: proneural cells (either proliferative PNs or quiescent PNq), persister cells (P), and mesenchymal cells (either proliferative MESs or quiescent, MESq). The cell dynamics between different compartments is summarized in Figure S4 (and in the Supplementary Information). It was assumed that PN cells may become MES cells, either directly, due to local vessel damage and hypoxia once the local cell density exceeds a critical threshold, or through a transient intermediate persister state induced by TMZ exposure. Both of these routes were associated with the emergence of TMZ resistance, as MES cells are assumed to be more resistant to TMZ than PN cells.

The methodology was first used to run simulations to explore the influence of the parameters on outcome, and to test the efficacy of the use of different dose spacings using murine parameters. Murine tumors were simulated without treatment (control), and treated with 3 TMZ doses, separated by 1, 4, 7 and 13 days. Human tumors were simulated without treatment, under standard TMZ therapy (6 cycles of TMZ given for 5 days, and resting periods of 3 weeks), and under two different courses of TMZ, increasing the spacing between doses. A first set of simulations was performed by increasing the rest periods between doses from 3 weeks to 9 weeks. Another set of computational studies was run by increasing the spacing between individual doses, while removing the rest periods. In particular, spacings of 8 and 12 days between doses were considered. Human and murine parameters were estimated from previous studies (see Table 1) and our own datasets. Virtual human simulations were fed with real patient data^35,36^ to generate realistic tumors *in silico*.

**Table 1:** Model parameters used to run murine and human simulations.

| PARAMETER | MEANING | VALUE<br>(MOUSE) | VALUE<br>(HUMAN) | UNITS | REFERENCE |
| --- | --- | --- | --- | --- | --- |
| $\tau_P$ | Division time<br>(proliferative<br>cells) | 19 | 48-72 | h | Estimated from<br>[29] |
| $\rho_M$ | Migration<br>coefficient | 0.12 | 0.4 | mm<sup>2</sup>/day | Estimated from<br>[29] |
| $\tau_D$ | Death time | 100 | 288-312 | h | Estimated from [29] |
| $\mu_{SQ}$ | Transition rate from proliferative state to quiescent | 0.3333-1 | 0.2-0.6 | day<sup>-1</sup> | Estimated from [29] |
| $\mu_{QS}$ | Transition rate from quiescent state to proliferative | 0.0166-0.1 | 0.03-0.2 | day<sup>-1</sup> | Estimated from [29] |
| $\mu_{SP}$ | Transition rate from PNs to persister | 0.0333 | 0.0333 | min<sup>-1</sup> | Estimated from [27] |
| $\mu_{PS}$ | Transition rate from persister to PNs | 0.2 | 0.2 | day<sup>-1</sup> | Explored in this work |
| $\mu_{PR}$ | Transition rate from persister to MESs | 0.0111 | 0.0111 | min<sup>-1</sup> | Estimated from [27] |
| $\mu_{PT}$ | Transition rate from PNs to MESs | 0.0208-0.0833 | 0.0208 | h<sup>-1</sup> | Explored in this work |
| $S_F$ | Survival fraction after TMZ dosing | 0.75 | 0.5 | unitless | Estimated from [38] |

The simulator was implemented in Julia (version 1.1.1). Simulation file processing and graphics were undertaken in MATLAB (R2021a, MathWorks). Simulations were performed on two 2.4 GHz, 16-core, 192 GB memory Mac Pro machines. Computational cost per murine simulation ranged from 5 to 10 minutes, while for humans, the computational cost ranged from 20 to 50 minutes per simulation.

## RESULTS

### Slow growth murine gliomas benefitted from increasing TMZ dose separation *in silico*

Virtual murine tumors were simulated as described in Methods, to explore the effect of different TMZ schemes in OS and MES content. A sustained increase in MES cell abundance was observed in untreated tumors (Figure 1A), reproducing the aforementioned PN-MES transition. These values are in agreement with real mouse data obtained in *in-vivo* experiments performed in this study. Notably, the tumor boundary was specially enriched in MES cells (Figure 1B).

**Fig. 1.**
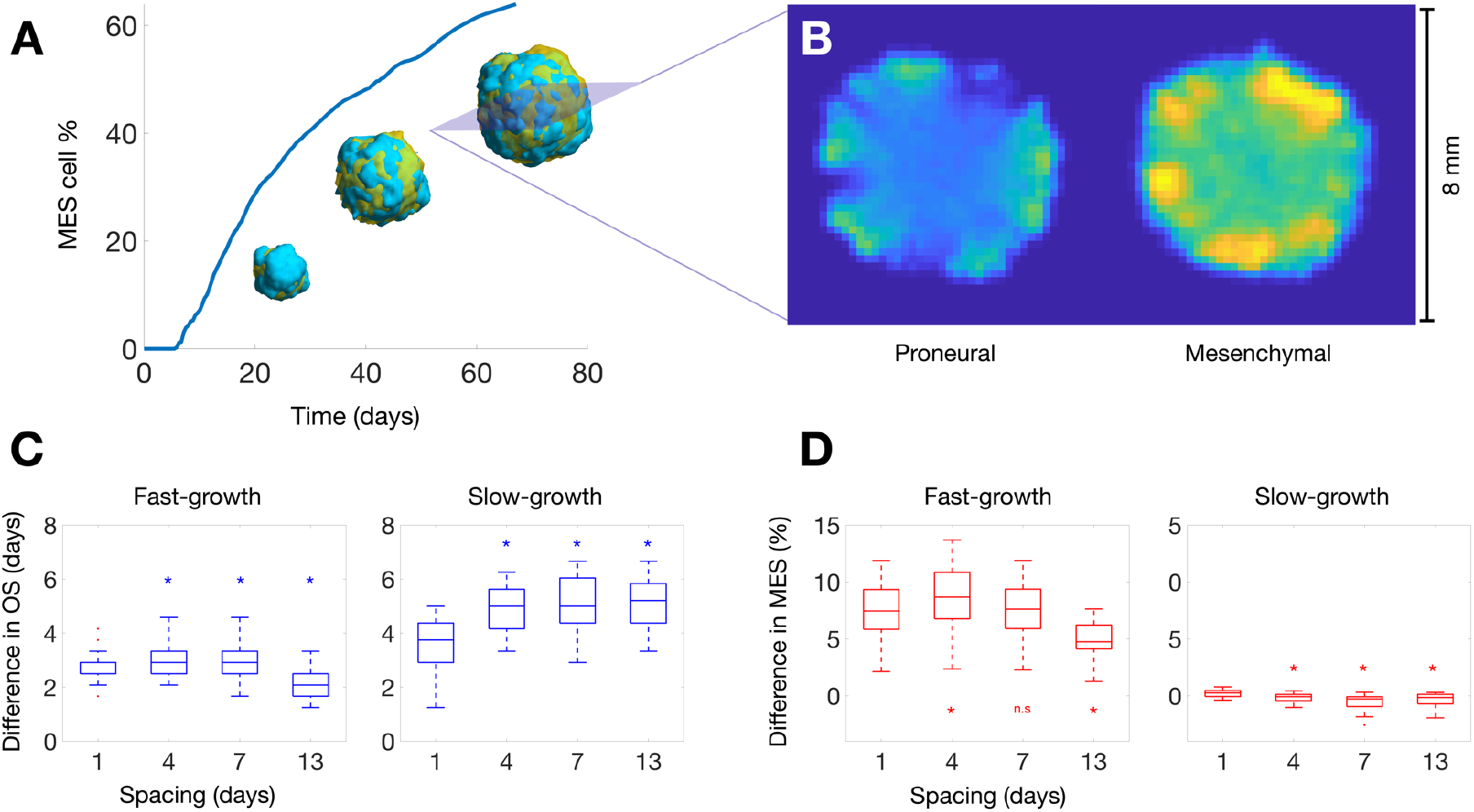
Mice tumor simulations predict an improvement in antitumor effect of TMZ in slow-growth GBMs. (**A**) Depiction of PN-MES transition in a single simulation, showing the evolution of MES cell abundance over time. 3D volumes are rendered at 50, 75 and 100% of simulation time (blue: PN cells; orange: MES cells). (**B**) Axial plane showing cell number per voxel in a simulated tumor at the end of simulation. Both PN and MES cell numbers belong to the same slice. (**C**) OS gain (in days) produced by TMZ dose spacing compared against control. 4-, 7- and 13-day spacings were compared against 1 day spacing (Wilcoxon signed-rank test). (**D**) MES cell increase (percentage) compared against control. 4-, 7- and 13-day spacings were compared against 1 day spacing (Wilcoxon signed-rank test, *P <= 0.05, n.s.=non-significant).

Treatment with TMZ yielded an increase in OS for both fast- and slow-growth tumors (Figure 1C). This survival increase was higher for slow-growth tumors; moreover, enlarging the spacing between doses produced a better response in this kind of tumors. Regarding resistance, TMZ induced a significant increase in MES cell content in fast-growth tumors (Figure 1D). However, slow-growth tumors did not undergo such increase; most of them remained with the same amount of MES cells as control tumors, or even reduced their MES levels. This effect was more evident for long spacings.

A robust observation was that virtual mice with slow-growth murine tumors had the largest survival increase when increasing the spacing between doses *in silico*. This benefit was preserved through the regions of the parameter space explored. Longer spacing between doses led to reduced mesenchymal component in the final tumors as compared to the 1-day spacing. Altogether, the simulations suggest that longer spacing between doses would be more effective against slow-growth gliomas, both in terms of OS and resistance development (Figures 1C-1D).

### Effect of protracted TMZ in two mouse glioma models with different proliferation kinetics

To validate the hypothesis that protracted TMZ regimes could offer therapeutic advantages depending on the degree of proliferation of glioma cells, mouse models were used, generated in our lab by overexpressing EGFRwt or EGFRvIII in p16/p19 ko subventricular zone (SVZ) progenitors. Both models generate gliomas in nude mice with a high penetrance and reproducibility. Notably, animals survive for two months after SVZ-EGFRwt cell intracranial injection, whereas SVZ-EGFRvIII tumors kill the animals much faster^31^, in agreement with the higher aggressiveness attributed to the mutated isoforms of EGFR^40,41^. Moreover, our previous analyses showed that tumors formed by SVZ-EGFRvIII cells were much more proliferative than those formed by SVZ-EGFRwt cells^31^.

Standard treatment of mouse glioma models with TMZ consisted of daily (5 days/week) i.p. injections of 10 mg/kg/day of the compound, which did not produce a survival benefit in animals bearing SVZ-EGFRwt or SVZ-EGFRvIII tumors (Figure 2A). Several TMZ regimes were then designed with three consecutive doses of three TMZ injections, separated by 1 day (X+1), 4 days (X+4), 7 days (X+7) or 13 days (X+13) (Figure 2B), although this last schedule could not be applied in the EGFRvIII model due to their faster growth. The graphs in Figure 2C show a clear reduction in SVZ-EGFRwt tumor growth in the X+7 and X+13 protracted schemes. Extending the interval between TMZ doses did not improve the response of SVZ-EGFRvIII bearing animals (Figure 2D). This result shows the truth (at least in mice) of the suggestion from the mathematical model that slowly-growing tumors are those that most benefit from protracted regimes.

**Fig. 2.**
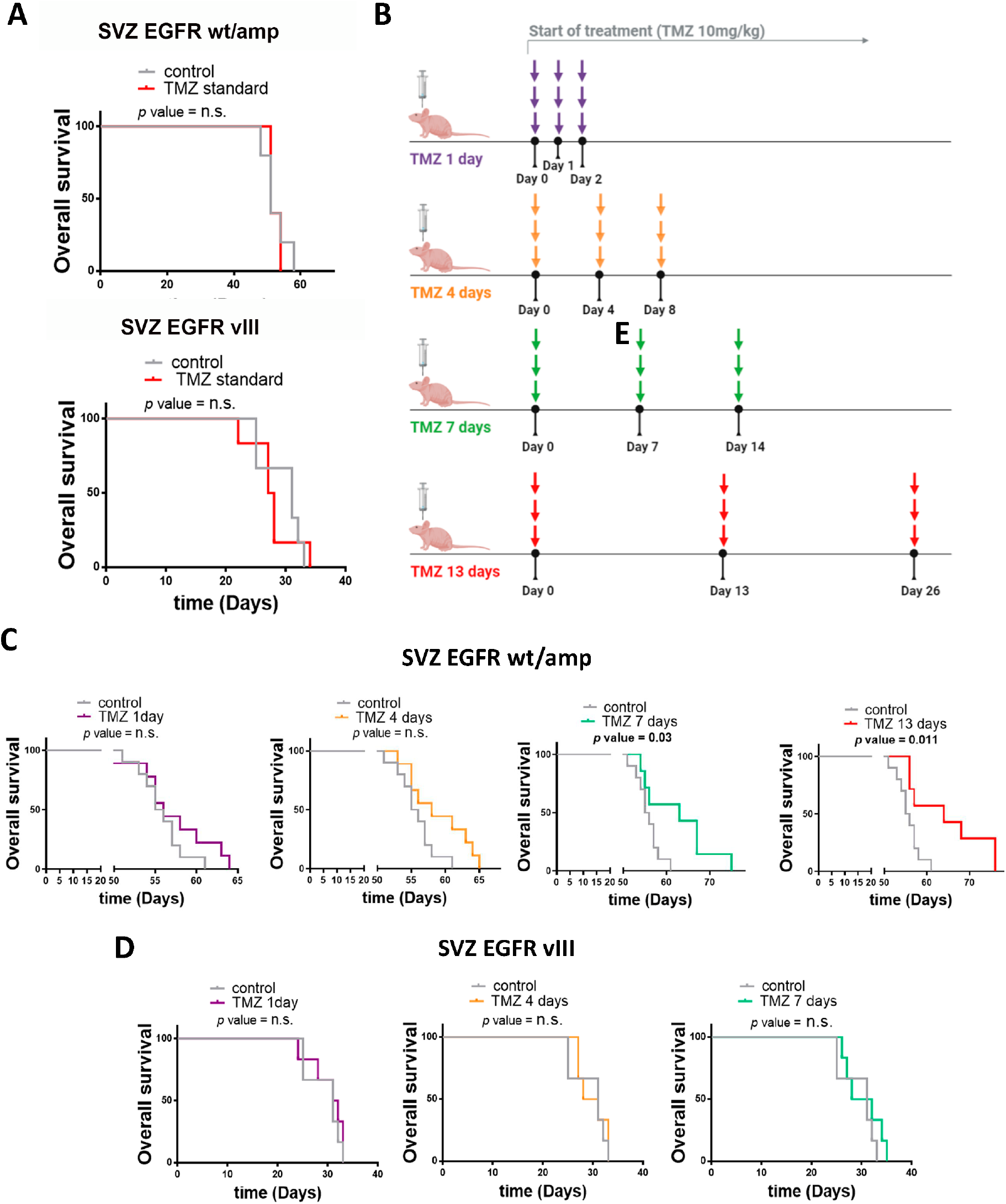
Increasing spacing between TMZ doses improves the anti-tumor effect of TMZ in the SVZ-EGFR wt/amp (lower growth rate) but not in the SVZ EGFR vIII (faster growth speed) model. (A) Kaplan-Meier overall survival curves of mice that were orthotopically injected with SVZ EGFR wt/amp (top) or SVZ EGFR vIII (bottom) cells and subsequently treated with intraperitoneal injections (five days per week) of temozolomide (TMZ) (10 mg/kg per day) (n = 6). (B) Representative scheme of the different TMZ schedules studied. (C-D) Kaplan-Meier overall survival curves of mice that were orthotopically injected with SVZ EGFR wt/amp (n = 9) (C) or SVZ EGFR vIII (n = 6) (D) cells and subsequently treated with intraperitoneal injections of TMZ (10 mg/kg per day) following the different treatment protocols explained in (B), each of them represented by its assigned color.

### Increasing spacing between TMZ doses does not increase cell death in the tumors but reduces the expression of persister-related genes

To understand the beneficial effect of PTS, tumor tissue in the SVZ-EGFRwt model was analyzed, comparing the control-treated with the X+4 (no response) and the X+13 (responsive) tumors. Notably, no changes were found in the tumor size in any of the schemes (Supplementary Figure 1A). In the X+13 scheme, a small decrease was observed in the number of proliferating cells (Supplementary Figure 1B), with no increase in the number of apoptotic cells (Figure 3A) compared to control tumors. However, in the X+4 scheme there was no change in proliferation, whereas an increase in the number of apoptotic cells was measured (Supplementary Figure 1C). These results suggest that the reduction in the tumor growth observed after X+13 protracted administration of TMZ is not mediated by an increase in the cell death response.

**Fig. 3.**
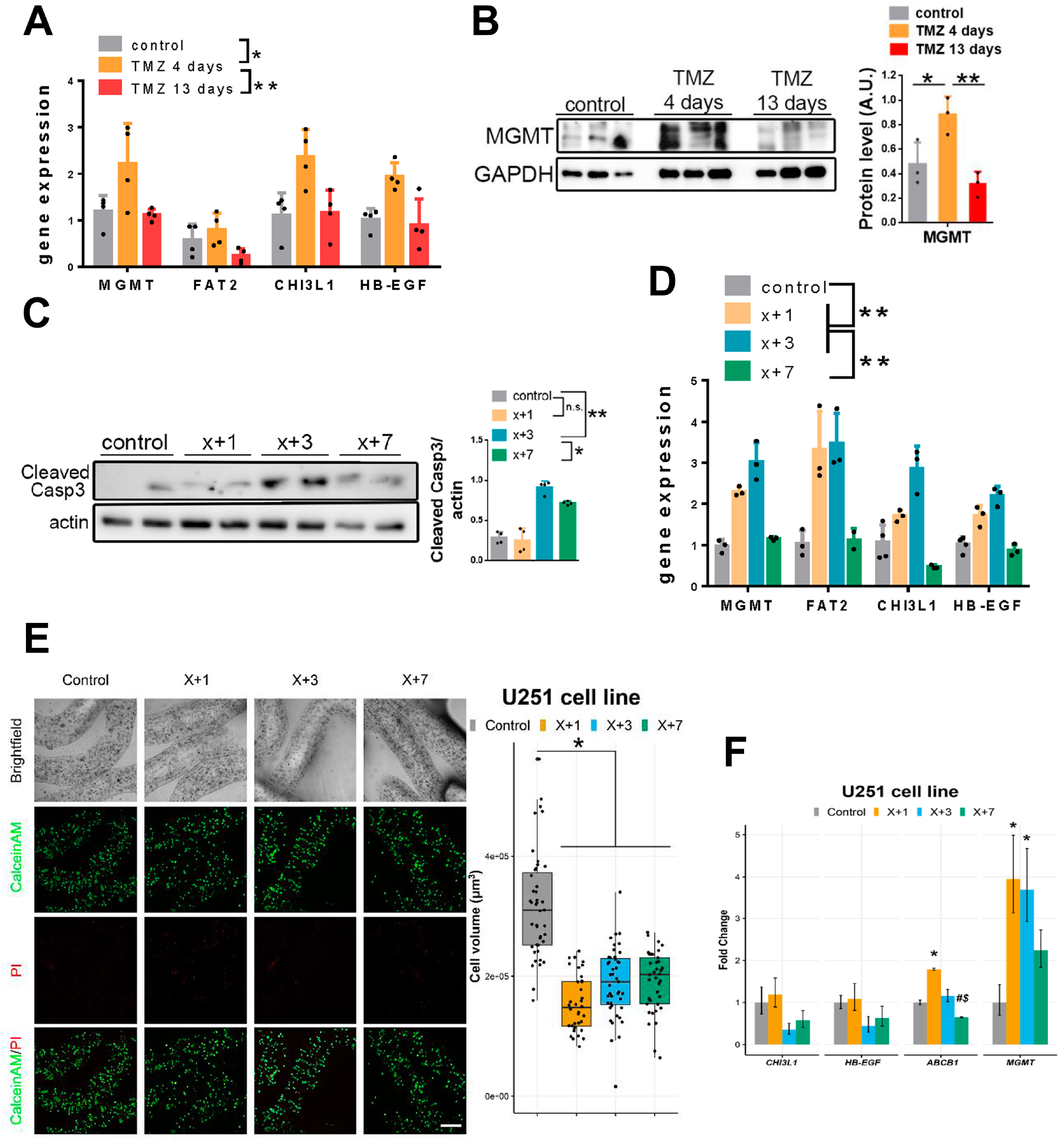
Increasing the spacing between TMZ treatments reduces the expression of persister-related genes *in vivo* and *in vitro.* (A) qRT-PCR analysis of persister-related genes in SVZ EGFR wt/amp tumors from (Fig. 2). *Actin* was used for normalization (n=4). (B) Western blot (WB) analysis and quantification of the expression of MGMT in SVZ tumors from (A). GAPDH was used for normalization. (C) WB analysis and quantification of Active Capase3 in SVZ EGFR wt/amp cells treated *in vitro* with different TMZ schedules: control, TMZ 1 day, TMZ 3 days and TMZ 7 days. *Actin* was used for normalization. (D) qRT-PCR analysis in SVZ EGFR wt/amp cells with the different treatment protocols. *Actin* was used for normalization (n=4). (E) Representative confocal microscopy images of U251 cells LIVE/DEAD labeled with CalceinAM/PI in alginate microfibers after 28 days when the treatments and cultivation of cells were completed. Quantification is shown as a box-plot on the right (n ≥ 3) Scale bar = 300 μm. (F) Relative gene expression levels of persister-related genes. *ACTB* was used for normalization (n = 3). *P ≤ 0.05, **P ≤ 0.01, n.s.=non significant. # indicates P<0.05 statistical difference compared to corresponding X+1 treatments. $ indicates P<0.05 statistical difference compared to corresponding X+3 treatments.

**Supplementary Fig. 1.**
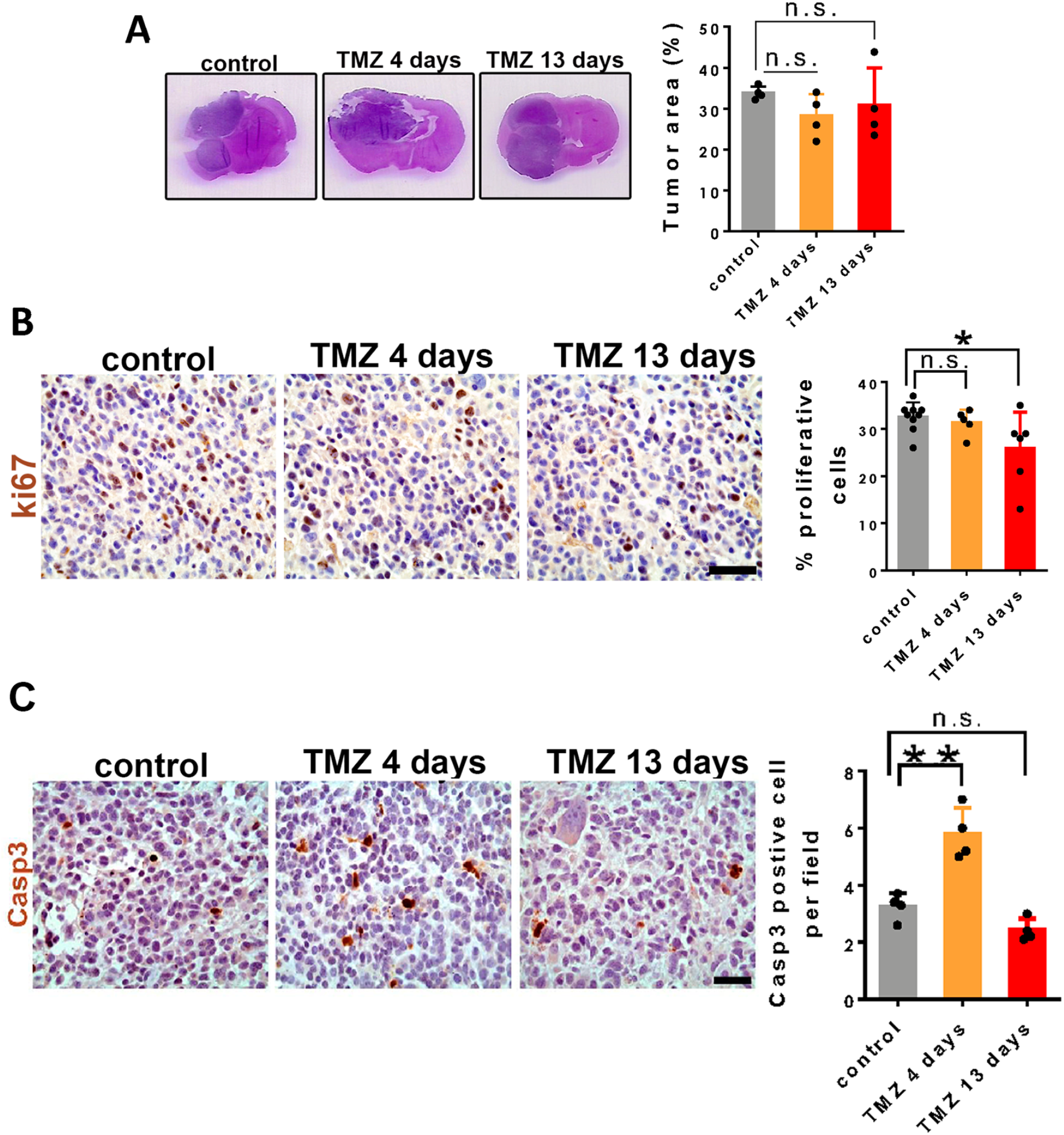
Effect of increasing spacing between TMZ doses in SVZ EGFR wt/amp tumors. (A) Representative histological images stained with hematoxylin and eosin (H&E) from coronal brain sections of SVZ EGFR wt/amp tumors from (Fig. 2) of control, TMZ 4 days and TMZ 13 days treatment condition. Quantification of tumor area percentage is shown on the right (n=3). (B) Representative pictures of immunohistochemical (IHC) staining of ki67 in the same tumors from (A) and quantification of the percentage of proliferative cells on the right (n=3). (C) Representative pictures of IHC staining of Active Caspase3 in SVZ EGFR wt/amp in the same tumors from (A). Quantification is shown on the right (n=3).*P ≤ 0.05, n.s.=non significant.

As previously mentioned, persister cells represent an intermediate phenotype arising before the development of TMZ resistance in gliomas^29^. To test whether different modalities of protracted TMZ treatment could affect the appearance of these population of cells, the expression of persister genes was analyzed. Notably, the expression of these markers was induced in tumors that had been treated with the X+4, but not with the X+13, regime, as compared to untreated animals (Figure 3A). Interestingly, one of these genes is *MGMT,* whose expression is strongly associated with TMZ resistance^42^. It was confirmed that MGMT protein was indeed being accumulated in the X+4 but not in the X+13 scheme (Figure 3B). These results suggest that extending the rest periods between TMZ treatments not only improved the anti-tumor effect of the drug, but also reduced the appearance of a persister state in the glioma cells.

In order to test whether the effect of protracted TMZ was cell-autonomous, SVZ-EGFRwt cells were treated with different schedules of TMZ *in vitro.* An increase was observed in caspase 3 cleavage in the X+3 scheme (Figure 3C), accompanied by the accumulation of the expression of persister genes, including *MGMT* (Figure 3D). Notably, an extended period between TMZ doses (X+7 scheme) did not produce a significant increase in Caspase 3 (Figure 3C) or in the expression of persister genes (Figure 3D). These results suggest that the anti-tumor mechanism of protracted TMZ doses is the same *in vivo* and *in vitro* and does not depend on the tumor microenvironment.

To confirm these results in human cells, U251 cells, grown in conventional 2D conditions, were exposed to three doses of TMZ (100 μM), comparing daily (X+1) and 3-day (X+3) schedules (Supplementary Figure 2A). Both regimes were able to reduce the growth of the cells compared to the control, according to the change in doubling time (Supplementary Figure 2B). However, in both cases an increase in the expression of resistance-related markers in the TMZ treated cells was detected (Supplementary Figure 2C). It was therefore determined to test the effect of PTS in conditions where the cells would have slower growth. For this purpose, U251 cells were immobilized in alginate microfibers and grown them in 3D conditions. The cells were then exposed to three doses of TMZ (100 μM), comparing daily (X+1), 3-day (X+3) and 7-day (X+7) schedules (Figure 3E). In all cases, TMZ reduced the growth of the cells (Figure 3F). Notably, X+7 decreased the expression of persisters-related genes (*CHI3L1, HB-EGF,* and *ABCB1*) in TMZ treated cells (Figure 3F). These results suggest that enlarging the intervals between doses could reduce the appearance of a persister phenotype *in vitro* in human glioma cells, at least in conditions of reduced proliferation.

**Supplementary Fig. 2.**
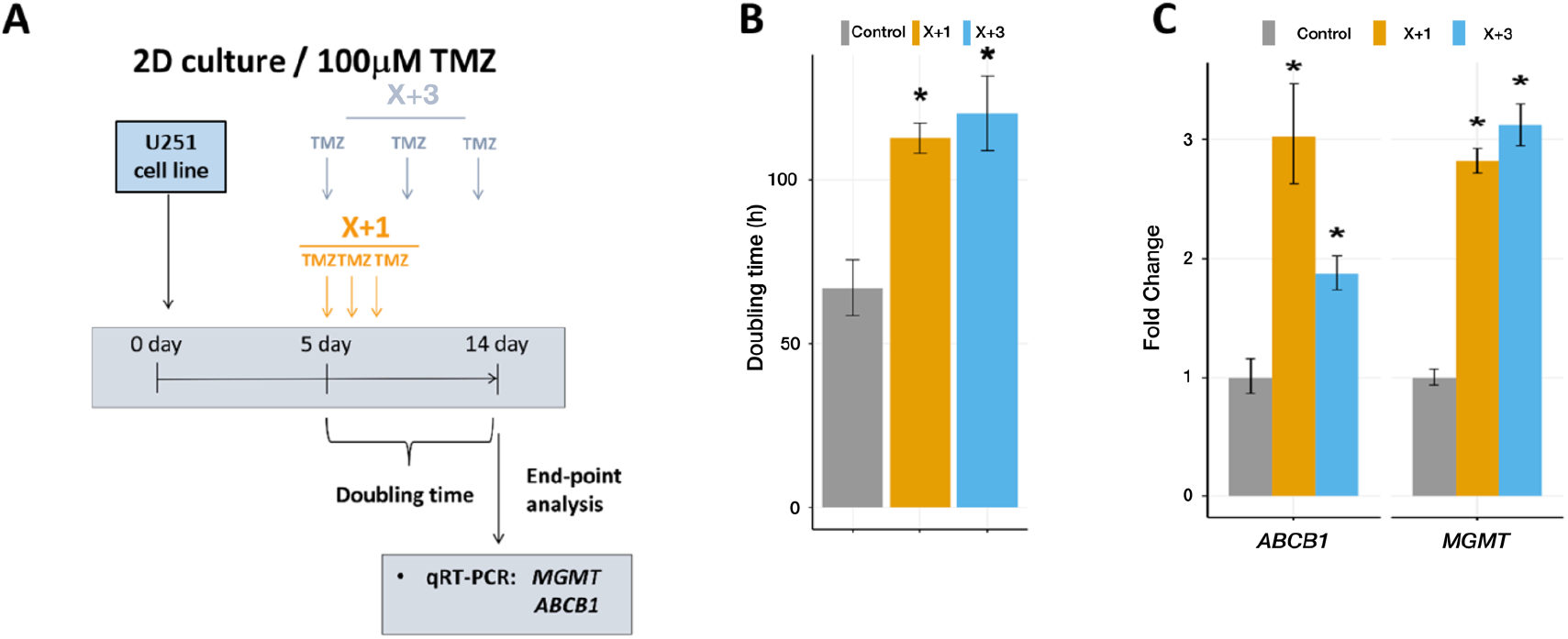
Effect of TMZ *in vitro* in 2D cultures. (A) TMZ treatment scheme in long-term 2D cell cultures of U251 glioblastoma cell line. (B) Doubling Time (dt) calculated using RTCA 1.2.1 software illustrates the anti-proliferative effects of TMZ treatments in U87 cell line. (C) Relative gene expression levels of *ABCB1* and *MGMT* in U251 cells obtained by qRT-PCR. All values are expressed as mean ± SD and *ACTB* was used for normalization (n = 3).

### Protracted TMZ induced a change in the phenotype of slowly-proliferating gliomas, mediated by epigenetic changes

Our previous characterization of the SVZ-EGFRwt model shows that these cells express MES features^30,31^. Interestingly, an *in-silico* analysis of two proliferation markers, PCNA and MKI67, was performed, and it was found that MES gliomas expressed the lowest levels of these genes (Supplementary Figure 3A), suggesting that tumors with this phenotype are less proliferative than the other two subtypes (classical (CL) and PN).

**Supplementary Fig. 3.**
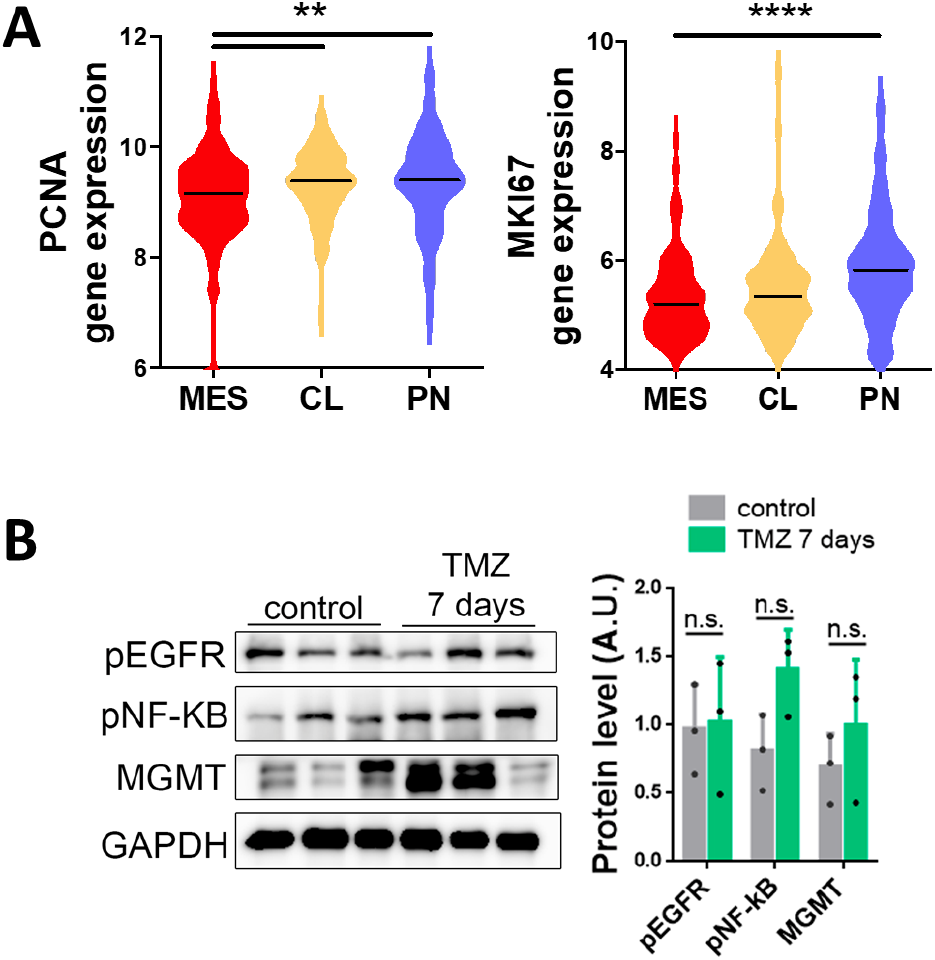
Expression of proliferation-related markers in different subgroups of gliomas. (A) RNA-seq analysis of *PCNA* (left) and *MKI67* (right) in a TCGA cohort stratified in to three groups: mesenchymal (MES), classical (CL) and proneural (PN) tumors. (B) Western blot analysis of phosphorylated EGFR (pEGFR) and NF-kB (p65) (pNF-kB) in SVZ-EGFRvIII tumors from Fig. 2. GADPH was used as a loading control. Quantification is shown on the right (n= 3).

It has previously been shown that the MES profile of SVZ-EGFRwt tumors depends on the activation of the EGFR/NFkB signaling pathway. Notably, EGFR has been associated with TMZ resistance in gliomas^43^. The activation status of this receptor in SVZ-EGFRwt tumors treated with TMZ was therefore explored, both in responsive and non-responsive schedules. A clear downregulation of the levels of phosphorylation of EGFR and NF-kB was observed in the X+13 tumors (Figure 4A). In tumors from the X+4 regime there was an increase in phospho-EGFR, although no significant changes were observed in the amount of phospho-NFkB (Figure 4A), a MES driver in gliomas^39^. The changes observed in the X+13 responsive tumors were paralleled by an increase in the expression of several PN markers (Figure 4B) and the downregulation of the transcription of MES genes (Figure 4C), compared to control tumors. These results suggest that less intensive TMZ schedules might be reducing the aggressiveness of gliomas by inducing a MES to PN phenotypic change. Notably, no changes in the amount of phosphorylation of EGFR or NF-kB were observed in TMZ-treated SVZ-EGFRvIII tumors compared to controls (Supplementary Figure 3A), reinforcing the lack of response of these gliomas to the drug.

**Fig. 4.**
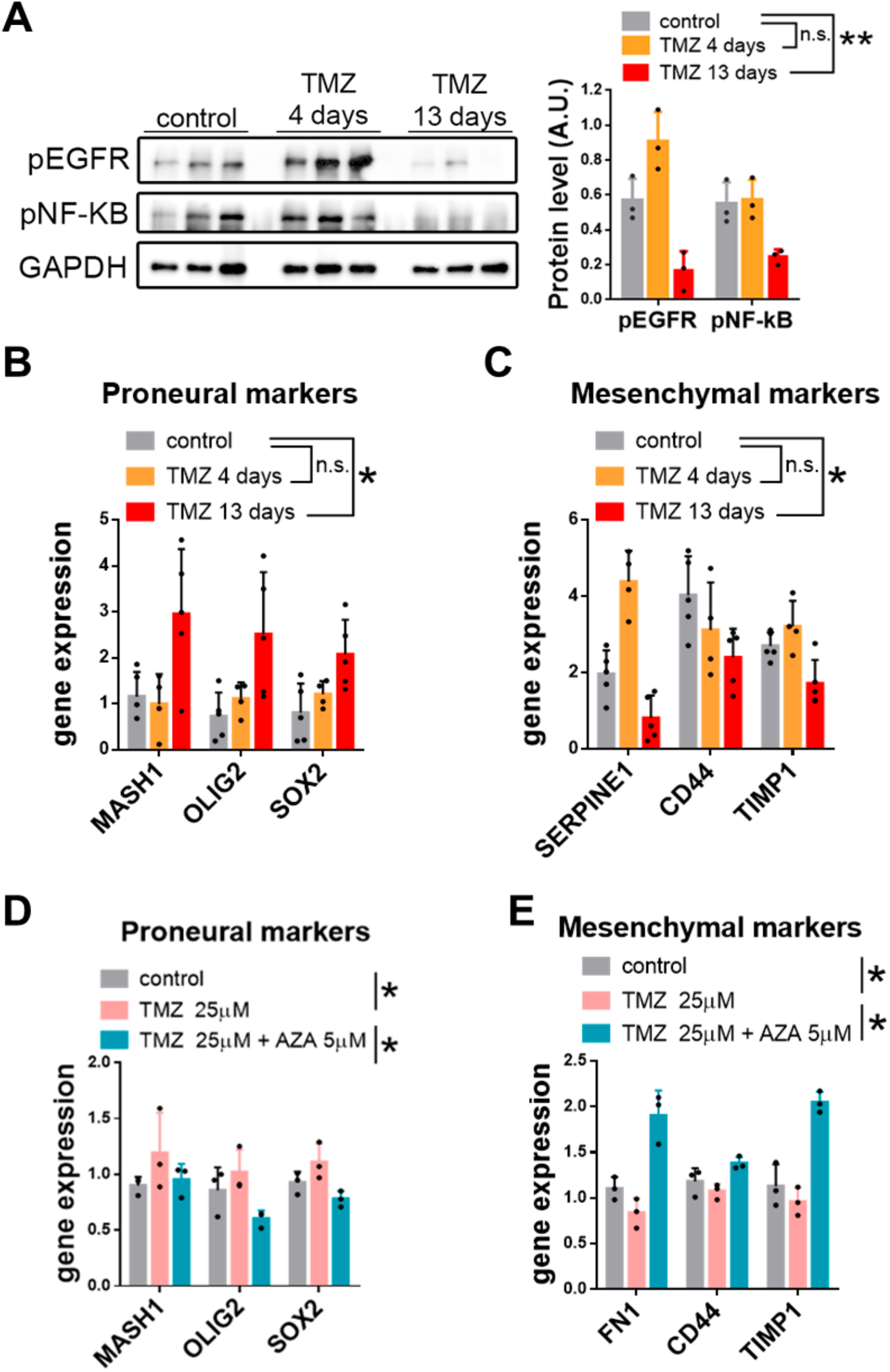
Mechanistic studies. Change of glioma phenotype for different TMZ treatment schedules. (A) Western blot analysis and quantification of phosphorylated EGFR (pEGFR) and NF-kB (p65) (pNF-kB) in SVZ EGFR wt/amp tumors from (Fig. 2). GADPH was used as a loading control (n= 3). (B-C) qRT-PCR analysis of Proneural (B) and Mesenchymal subtype (C) related genes in SVZ EGFR wt/amp tumors from (A). *Actin* was used for normalization (n=4). (D-E) Analysis of the expression of Proneural (D) and Mesenchymal markers (E) transcription by qRT-PCR in SVZ EGFR wt/amp cells cultured in the presence of 25 μM TMZ, with or without azacytidine (AZA) (5 μM). *Actin* was used for normalization (n= 3). *P ≤ 0.05, **P ≤ 0.01, n.s.=non significant.

To study the anti-tumor mechanisms of TMZ more deeply, SVZ-EGFRwt cells were incubated *in vitro* in the presence of TMZ (25 μM) for 8 days. As noticed in tumors treated with TMZ, an increase in the expression of PN markers was observed (Figure 4D). Notably, this effect was reverted in the presence of the DNA-methyltransferase inhibitor 5-aza-2’deoxycytidine (AZA) (Figure 4D), suggesting that TMZ might be inducing the expression of these genes by a shift in the DNA methylation pattern, previously proposed as a mechanism of action of this drug in glioma cells^44^. A decrease in the expression of MES genes was also noticed, which did not occur in the presence of AZA (Figure 4E). Therefore, it was hypothesized that the anti-tumor effect of protracted schemes of TMZ might be mediated, at least in part, by a MES-to-PN transition of the tumor cells induced by epigenetic changes, the opposite of what would normally happen during tumor progression^45^. This would increase the effect of giving more time to persister cells to revert their phenotypes to the PN phenotype.

### TMZ dose can be increased in a protracted scheme to enhance the anti-tumor effect and reduce toxicity

One of the potential benefits of long spacing between TMZ cycles could be a reduction in the toxicity of the drug, which may perhaps allow the CT dose to be increased. To test this hypothesis, the amount of TMZ administered to the animals after the intracranial injection of SVZ-EGFRwt cells was increased to 50mg/Kg/day. The same schemes for TMZ treatment were used as in Figure 2, and confirmed that the longest period between cycles was the most effective in reducing tumor growth (Figure 5A). The day after the last TMZ cycle, blood was collected from the animals to perform a white-cell count. One of the most common adverse effects of chemotherapy with TMZ is myelosuppression, including thrombocytopenia and leukopenia^46^. Indeed, a reduction was observed in all the numbers in the X+1 regime that reached a statistically significant value for the decrease in thrombocytes (Figure 5B). Notably, extending the period between doses was able to revert to normal the leukocyte and thrombocyte counts (Figure 5B), indicating that the toxicity of TMZ was reduced, even though the anti-tumor effect was increased in comparison to the lower dose regime.

**Fig. 5.**
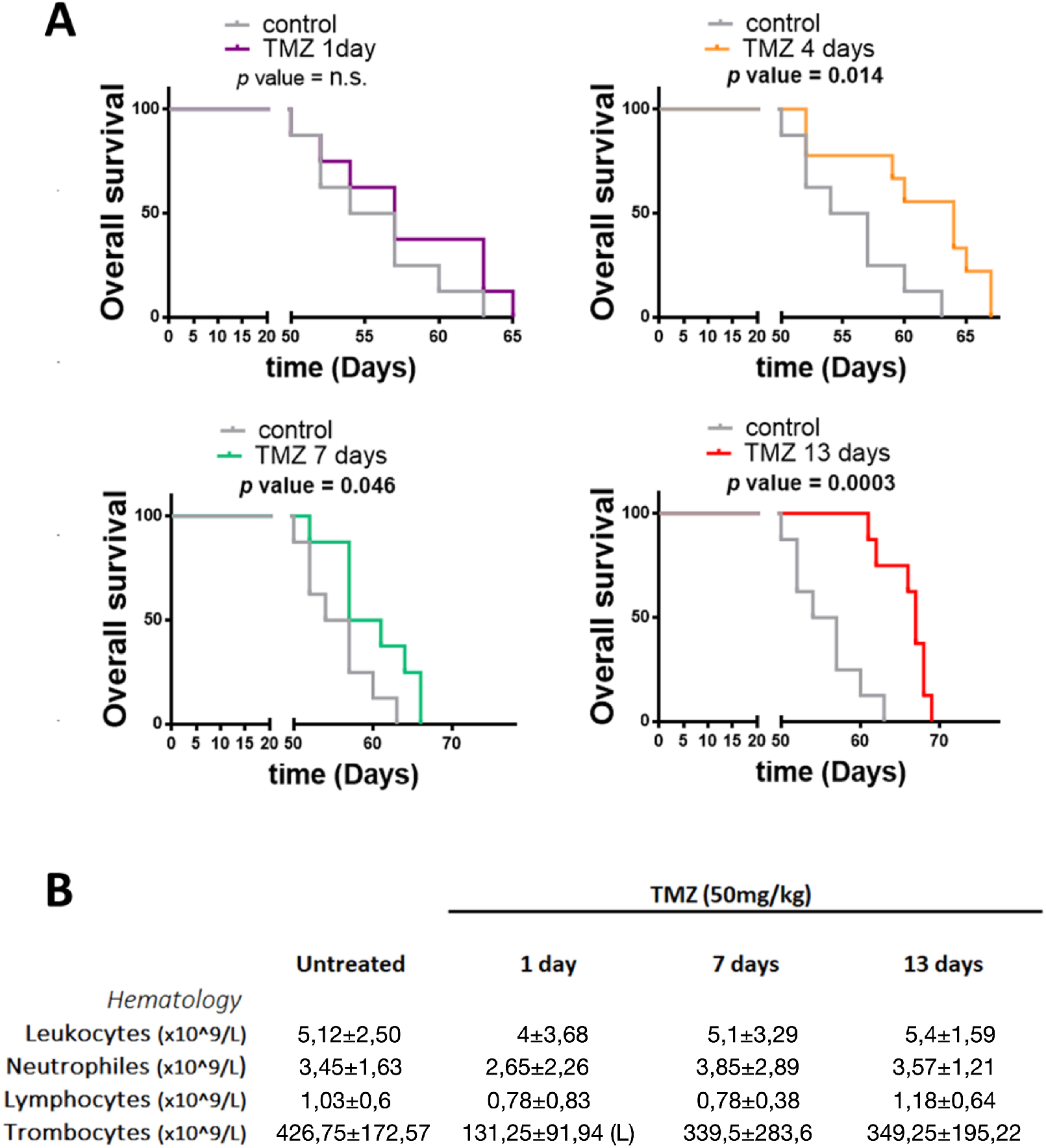
Reduced toxicity of protracted TMZ schemes. (A) Kaplan-Meier overall survival curves of mice that were orthotopically injected with SVZ EGFR wt/amp cells and subsequently treated with intraperitoneal injections of a higher dose of TMZ (50 mg/kg per day) following the different TMZ protracted schemes (n = 9). (B) Hematology results in mouse blood samples from (A).

### Virtual clinical trials suggest how to translate the results in experimental models to human patients

To extend previous *in vivo* findings, many sets of virtual clinical trials were performed, based on the mathematical model with human parameters. The potential effect of PTS on tumor growth dynamics was assessed by (i) enlarging the rest periods between cycles, and (ii) testing long separation times between individual doses without rest periods (Figure 6). Simulated tumors were separated in slowly-growing and fast-growing GBMs. All *in-silico* patients were given 30, 60 or 90 doses of TMZ 7 weeks after diagnosis (depending on therapy regime).

**Fig. 6.**
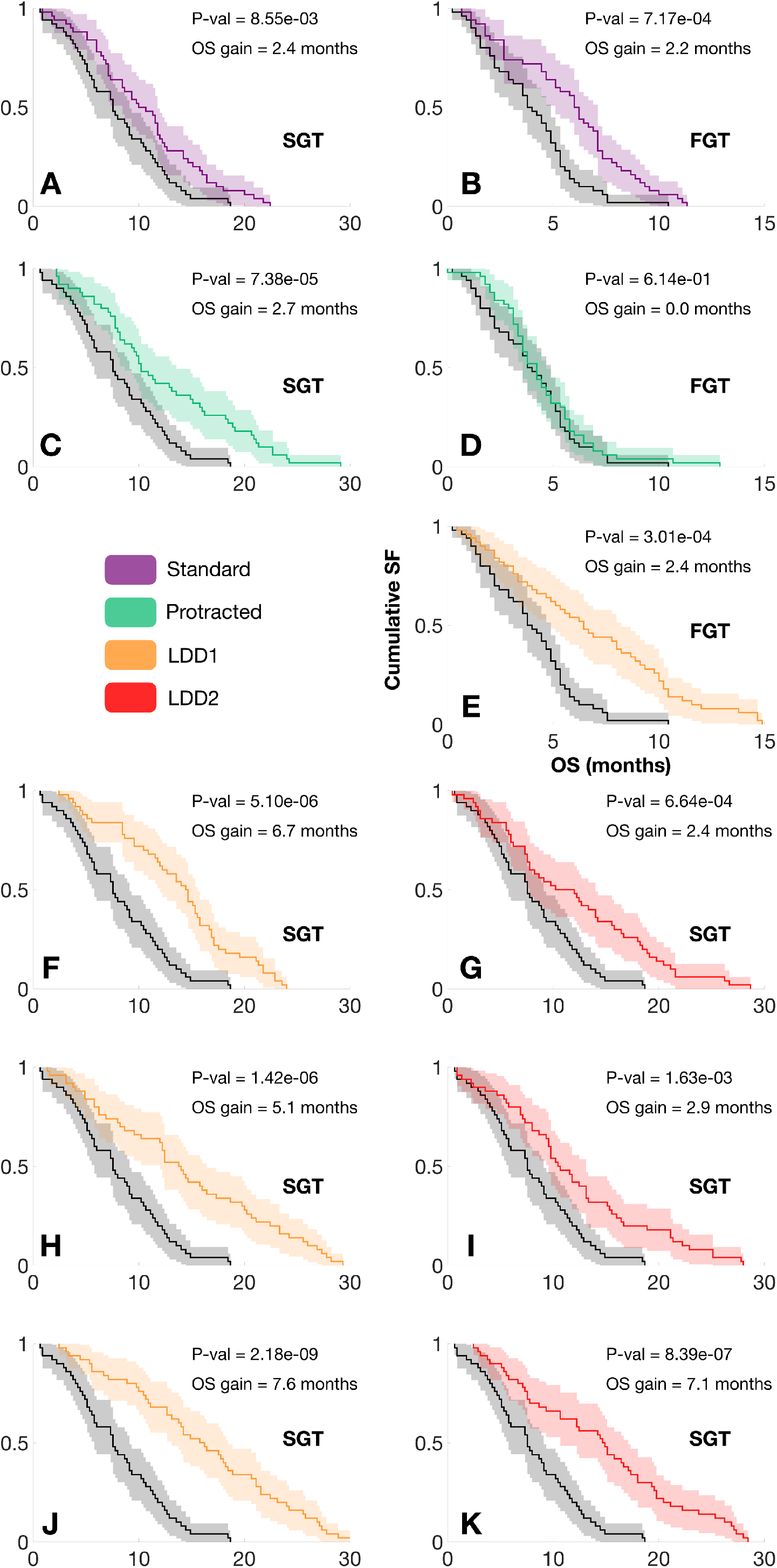
Kaplan-Meier survival curves for PTS *in silico.* Standard therapy (red) consists of 6 cycles of TMZ, with 5 doses per cycle, and 23 days of resting period. Protracted therapy (green) consisted of standard therapy, with a resting period of 9 weeks. Therapies with long spacing between doses 1 (LDD1, orange) correspond to doses separated by 8 days without any rest periods. Therapies with long spacing between doses 2 (LDD2, red) consisted of individual doses spaced by 12 days without rest periods. **A)** Slow-growing simulated tumors with 6 cycles of standard therapy. **B)** Fast-growing simulated tumors with 6 cycles of standard therapy. **C)** Slow-growing simulated tumors with 6 cycles of protracted therapy. **D)** Fast-growing simulated tumors with 6 cycles of protracted therapy. **E)** Fast-growing simulated tumors with 6 cycles of LDD1 therapy. **F)** Slow-growing simulated tumors with 6 cycles of LDD1 therapy. **G)** Slow-growing simulated tumors with 6 cycles of LDD2 therapy. **H)** Slow-growing simulated tumors with 12 cycles of LDD1 therapy. **I)** Slow-growing simulated tumors with 18 cycles of LDD1 therapy. **J)** Slow-growing simulated tumors with 12 cycles of LDD2 therapy. **K)** Slow-growing simulated tumors with 18 cycles of LDD2 therapy.

Standard TMZ schemes (5 consecutive doses, rest period of 3 weeks) showed a beneficial effect in terms of survival for both tumor types (Fig. 6A, 6B), with a median survival difference of nearly 2 months, in line with clinical experience^12^. Increasing the rest period to 9 weeks improved survival for slow-growth tumors, with an increased number of longer survivors (Fig. 6C). Fast-growing tumors did not benefit from the increase in spacing between doses (Fig. 6D), in line with our observations in animal models.

On the other hand, all trial branches benefited from increasing time intervals between doses (without resting periods), both for slow and fast-growing tumors. As the number of cycles given was increased, differences in median survival increased remarkably.

## DISCUSSION

TMZ is the standard of care for newly diagnosed GBM, but the effect of this alkylating agent is schedule-dependent^47^. Genetic or acquired resistances to TMZ can easily develop, and a strict regimen must be followed for a favorable result to be obtained^48,49^. The design of cytotoxic chemotherapy (CT) and radiation therapy schedules is typically based on the basic principle of delivering the maximum tolerated dose in the minimum time possible (MTDMT). The rationale behind this is to avoid potential tumor repopulation during treatment, or in the periods without treatment, and thus to achieve low tumor-cell numbers compatible with patient cure. Although this is certainly the way to go when CT is intended as a curative treatment, it is not obvious that it would be the best strategy when it is known that treatment can only control disease for a limited time.

Intensification of TMZ delivery schemes has been studied for either newly diagnosed^31^ or recurrent^32,33^ high-grade gliomas without positive results on OS and with increased toxicity^38^. However, no previous studies have considered effective dose-reduction schemes with longer time spacing between treatments. Different studies based on mathematical models have argued that cytotoxic therapies with larger time intervals between doses could provide survival benefits. However, these mathematical models are based on saturable growth models, where tumors proliferate less on average as they grow larger, and the phenomenon is lost when exponential growth models are considered^21^. The situation would be worse in the context of super-exponential growth models^22^ where the opposite result would be obtained, i.e. survival benefits from the MTDMT approach.

This work studies, *in vitro, in vivo* and *in silico* using mathematical models, the main factor limiting the effectiveness of treatment in gliomas: the development of resistance. Resistance acquisition in the mathematical model was assumed to be due to two biological facts: PN-MES transition, and persister induction by TMZ. Accounting for resistances in our stochastic mesoscopic discrete simulation model led to improved survival and reduced resistance when increasing the interval between doses in virtual mice. The reduction of resistant cells stemmed from the fact that longer spacing between doses allowed persister cells to revert their phenotypes to PN. Clearly, when persistence time is longer than the spacing between doses, persister cells receive additional TMZ doses and the emergence of resistance is triggered. Very interestingly, for slowly-growing tumors, as TMZ kills a fraction of both PN and MES cells, the resistance level at the end of the simulation was observed to be smaller than its control counterpart, as the spontaneous PN-MES transition is being reduced due to TMZ killing PN cells. Thus, increasing the spacing between doses provides the same effect as a reverse MES-PN transition. This setup was translated to the real world, both in *in vitro* and *in vivo*. Results in mice were in accordance with model suggestions: slow-growing tumors benefited most from PTS. In fact, fast-growing tumors did not show an improvement in OS, as expected.

A phenotypic change in slowly-growing tumors subject to increased spacing between doses was observed *in vivo*, with a reduction in the levels of phosphorylated EGFR and NF-κB, s associated with a MES-to-PN switch, as recently shown^30^. This transition could explain the reduced aggressiveness of the tumors after long-cycle TMZ treatment, which does not seem to depend on changes in proliferation and/or survival of tumor cells. Notably, our data suggest that this MES to PN switch does not depend on the tumor microenvironment and can be reverted in the presence of AZA, the known epigenetic regulator. Changes in DNA methylation has already been associated with the response to TMZ in a time and dose-dependent manner^44^. Moreover, it has been previously shown that the persister state is also linked with alterations in the levels of histone acetylation and with chromatin remodeling processes^29^. Therefore, it could be hypothesized that epigenetic changes might be responsible for the appearance of resistances, but also for some of the anti-tumor effects of TMZ, all linked to alterations in the transcriptomic profiles of GBMs. Furthermore, our results might explain why extensive TMZ treatment did not alter the survival of PN gliomas, but was beneficial for the more aggressive MES subtype^50^. Anyhow, these results not only emphasize the potential clinical relevance of PTS for slowly-growing GBMs, but also indicate that the biological assumptions taken in the model of action of this compound are not far-fetched, and should be explored in deeper detail to keep improving our knowledge about GBMs and the best way to treat them.

The experimental evidence shown here supports the fact that PTS could be beneficial for GBM patients in terms of survival, resistance and toxicity so far. To gain some insight into the improvements that could be expected in human patients, many virtual trials were performed based on the mathematical model, to explore the consequences PTS in clinical scenarios. The aim was to provide a broad exploration of the outcomes of protracted regimes, and a proof of concept that, if taken with caution, may be helpful in guiding the design of future clinical trials.

The output yield by the virtual clinical trial was in agreement with the experimental results obtained in this work. Standard TMZ therapy showed a moderate improvement in terms of survival, in line with clinical experience. Fast-growing tumors did not benefit from increasing the rest periods between cycles, but they did benefit from enlarging the spacing between doses. Slow-growing tumors benefitted not only from every alternative therapy scheme, but also from an increase in the number of cycles given due to the reduction in the MES component, and thus in tumor resistance. This points to alternative schemes that would allow for more TMZ doses to be given, while keeping resistances stable and with lower toxicity. There were several interesting implications from these virtual trial studies. The first was that extending the rest period between 5-dose cycles from 3 to 9 weeks showed a significant improvement in survival for patients with slow-growing GBMs, suggesting an easy-to-apply upgrade in the standard of care. The second was that there may be room for optimizing TMZ schedules in GBMs, as the best improvements in survival came from schemes without rest periods, and 8-12 days between doses. Due to their lower dose density, these schemes could be less harmful in terms of toxicity and, following both virtual and experimental results, a reduced resistance should also be expected, making them an alternative option for clinical implementation.

In conclusion, our combination of *in-silico* simulations, *in-vitro* and *in-vivo* studies showed that TMZ administration schedules with increased time spacing between doses may reduce toxicity, delay the appearance of resistances and lead to survival benefits mediated by changes in the tumor phenotype, which was especially important for slowly growing gliomas. The experimental results were extended to human patients showing different ways to improve survival based on the same concept.

## ACKNOWLEDGMENTS

This work has been supported by the James S. Mc. Donnell Foundation (USA) 21st Century Science Initiative in Mathematical and Complex Systems Approaches for Brain Cancer (Collaborative award 220020560); Ministry of Education, Science and Technological Development, Republic of Serbia (ref. number 451-03-9/2021-14/200007); Ministerio de Ciencia e Innovación, Spain (grant number PID2019-110895RB-I00). JJ-S is supported by the University of Castilla-La Mancha under fellowship grant number 2020-PREDUCLM-15634. We thank Luis Enrique Ayala-Hernández and Juan Belmonte-Beitia for their thoughtful discussions regarding continuous mathematical descriptions of GBM growth and response to TMZ, Julián Pérez-Beteta for providing us real patient data required to parametrize the discrete model, as well as Sofija Jovanović Stojanov and Ana Podolski-Renić for their valuable assistance in performing and analyzing results in human glioblastoma 2D and 3D cell culture.

---

**Supplementary Fig. S4.**
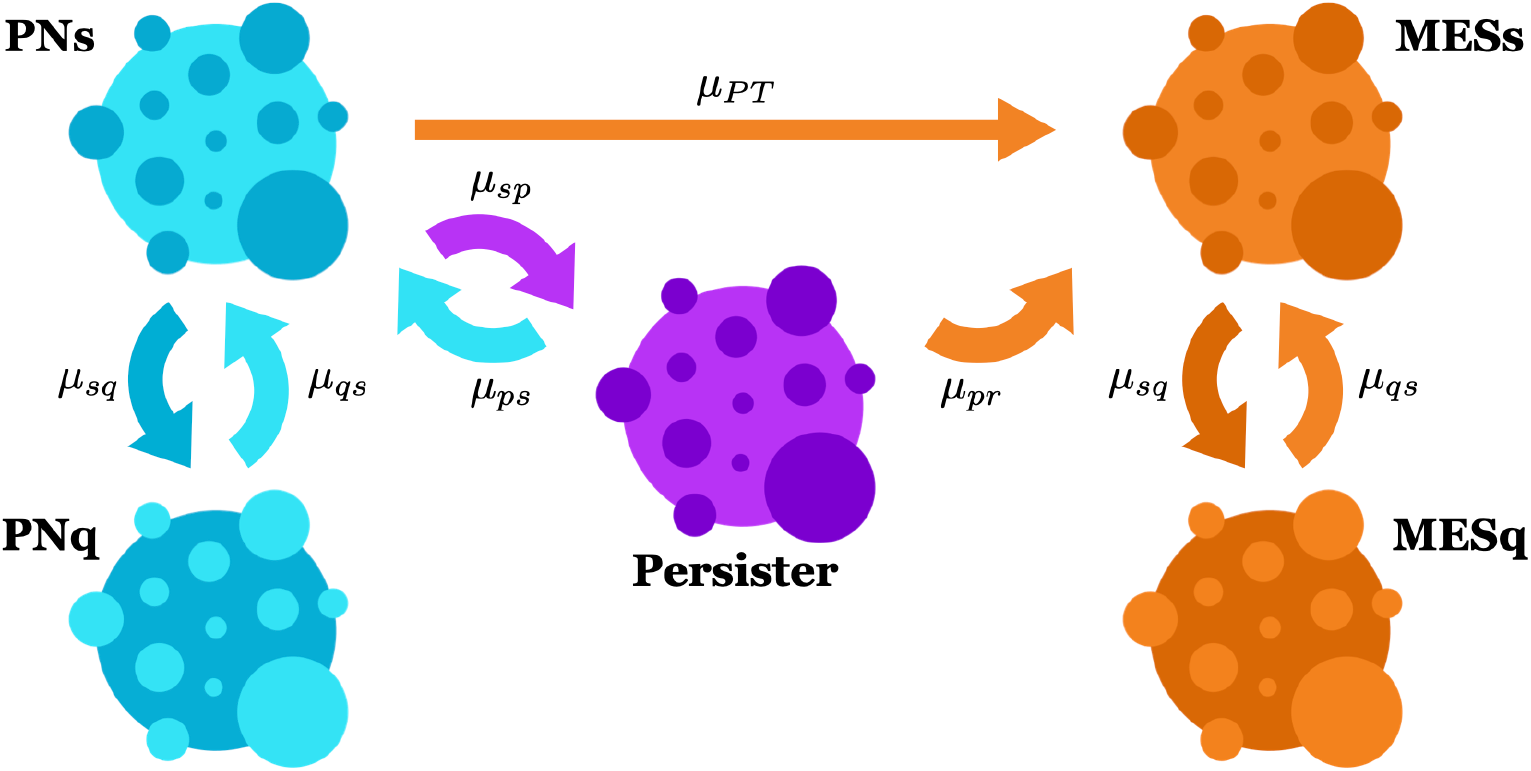
Allowed cell transitions in the model. Both PN and MES cells share the same proliferative-quiescent dynamics. PNs can become MES either directly or due to TMZ exposure. In the latter case, the transition occurs through a transient reversible persister state.

**Supplementary Fig. S5.**
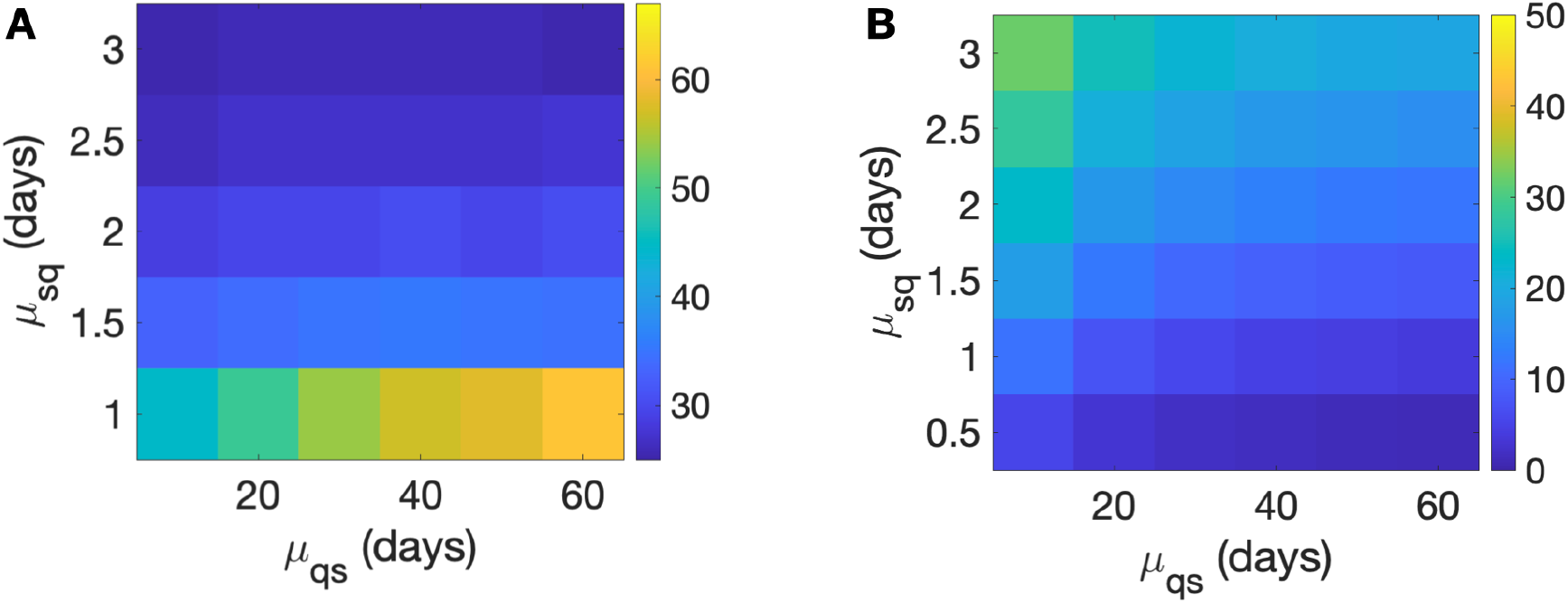
Overall survival in days (**A**) and Ki67 % (**B**) obtained from different combinations of initial ranges of parameters μ_sq_ and μ_qs_.

## REFERENCES

1. Brat DJ, Aldape K, Colman H, et al. cIMPACT-NOW update 3: recommended diagnostic criteria for “Diffuse astrocytic glioma, IDH-wildtype, with molecular features of glioblastoma, WHO grade IV”. Acta Neuropathol. 2018; 136(5):805–810.

2. Louis DN, Perry A, Wesseling P, et al. The 2021 WHO Classification of Tumors of the Central Nervous System: a summary. Neuro Oncol. 2021;23(8):1231–1251.

3. Louis DN, Perry A, Reifenberger G, et al. The 2016 World Health Organization Classification of Tumors of the Central Nervous System: a summary. Acta Neuropathol. 2016; 131(6):803–820.

4. Louis DN, Wesseling P, Aldape K, et al. cIMPACT-NOW update 6: new entity and diagnostic principle recommendations of the cIMPACT-Utrecht meeting on future CNS tumor classification and grading. Brain Pathol. 2020; 30(4):844–856.

5. Brennan CW, Verhaak RG, McKenna A, et al. The somatic genomic landscape of glioblastoma. Cell. 2013; 155(2):462–477.

6. Weller M, van den Bent M, Preusser M, et al. EANO guidelines on the diagnosis and treatment of diffuse gliomas of adulthood. Nat Rev Clin Oncol. 2021; 18(3):170–186.

7. Stupp R, Mason WP, van den Bent MJ, et al. Radiotherapy plus concomitant and adjuvant temozolomide for glioblastoma. N Engl J Med. 2005; 352(10):987–996.

8. Zhang J, Stevens MF, Bradshaw TD. Temozolomide: mechanisms of action, repair and resistance. Curr Mol Pharmacol. 2012; 5(1):102–114.

9. Mur P, Rodríguez de Lope Á, Díaz-Crespo FJ, et al. Impact on prognosis of the regional distribution of MGMT methylation with respect to the CpG island methylator phenotype and age in glioma patients. J Neurooncol. 2015; 122(3):441–450.

10. Bady P, Delorenzi M, Hegi ME. Sensitivity Analysis of the MGMT-STP27 Model and Impact of Genetic and Epigenetic Context to Predict the MGMT Methylation Status in Gliomas and Other Tumors. J Mol Diagn. 2016; 18(3):350–361.

11. Wick W, Platten M, Weller M. New (alternative) temozolomide regimens for the treatment of glioma. Neuro Oncol. 2009; 11(1):69–79.

12. Gilbert MR, Wang M, Aldape KD, et al. Dose-dense temozolomide for newly diagnosed glioblastoma: a randomized phase III clinical trial. J Clin Oncol. 2013; 31(32):4085–4091.

13. Berrocal A, Perez Segura P, Gil M, et al. Extended-schedule dose-dense temozolomide in refractory gliomas. J Neurooncol. 2010; 96(3):417–422.

14. Taal W, Segers-van Rijn JM, Kros JM, et al. Dose dense 1 week on/1 week off temozolomide in recurrent glioma: a retrospective study. J Neurooncol. 2012; 108(1):195–200.

15. Nagane M. Dose-dense Temozolomide: Is It Still Promising?. Neurol Med Chir (Tokyo). 2015; 55 Suppl 1:38–49.

16. Wei W, Chen X, Ma X, Wang D, Guo Z. The efficacy and safety of various dose-dense regimens of temozolomide for recurrent high-grade glioma: a systematic review with meta-analysis. J Neurooncol. 2015; 125(2):339–349.

17. Pérez-García VM, Ayala-Hernández LE, Belmonte-Beitia J, et al. Computational design of improved standardized chemotherapy protocols for grade II oligodendrogliomas. PLoS Comput Biol. 2019; 15(7):e1006778.

18. Altrock PM, Liu LL, Michor F. The mathematics of cancer: integrating quantitative models. Nat Rev Cancer. 2015; 15(12):730–745.

19. Ribba B, Kaloshi G, Peyre M, et al. A tumor growth inhibition model for low-grade glioma treated with chemotherapy or radio-therapy. Clin Cancer Res 2012; 15:5071–5080

20. Mazzocco P, Honnorat J, Ducray F, Ribba B. Increasing the Time Interval between PCV Chemotherapy Cycles as a Strategy to Improve Duration of Response in Low-Grade Gliomas: Results from a Model-Based Clinical Trial Simulation. Comp Math Meth in Medicine 2015; 2015:297903.

21. Henares-Molina A, Benzekry S, Lara PC, García-Rojo M, Pérez-García VM, Martínez-González A. Non-standard radiotherapy fractionations delay the time to malignant transformation of low-grade gliomas. PLoS One. 2017; 12(6):e0178552.

22. Fisher BJ, Naumova E, Leighton CC, et al. Ki-67: a prognostic factor for low-grade glioma? Int J Radiat Oncol Biol Phys. 2002; 52(4):996–1001.

23. Liang J, Lv X, Lu C, et al. Prognostic factors of patients with Gliomas - an analysis on 335 patients with Glioblastoma and other forms of Gliomas. BMC Cancer. 2020; 20(1):35.

24. Perez-Garcia VM, Perez-Romasanta LA. Extreme protraction for low-grade gliomas: theoretical proof of concept of a novel therapeutical strategy. Math Med Biol. 2016; 33(3):253–271.

25. Fedele M, Cerchia L, Pegoraro S, Sgarra R, Manfioletti G. Proneural-Mesenchymal Transition: Phenotypic plasticity to acquire multitherapy resistance in glioblastoma. Int J Mol Sci. 2019; 20(11):2746.

26. Minata M, Audia A, Shi J, et al. Phenotypic plasticity of invasive edge glioma stem-like cells in response to ionizing radiation. Cell Reports 2019; 26:1893–1905.

27. Sharma SV, Lee DY, Li B, et al. A chromatin-mediated reversible drug-tolerant state in cancer cell subpopulations. Cell. 2010; 141(1):69–80.

28. Cabanos HF, Hata AN. Emerging Insights into Targeted Therapy-Tolerant Persister Cells in Cancer. Cancers (Basel). 2021; 13(11).

29. Rabé M, Dumont S, Álvarez-Arenas A, et al. Identification of a transient state during the acquisition of temozolomide resistance in glioblastoma. Cell Death Dis. 2020; 11(1):19.

30. Gargini R, Segura-Collar B, Herranz B, et al. The IDH-TAU-EGFR triad defines the neovascular landscape of diffuse gliomas. Sci Transl Med. 2020; 12(527).

31. Segura-Collar B, Garranzo-Asensio M, Herranz B, et al. Tumor-derived pericytes driven by EGFR mutations govern the vascular and immune microenvironment of gliomas. Cancer Res. 2021.

32. Dragoj M, Stojkovska J, Stanković T, et al. Development and Validation of a Long-Term 3D Glioblastoma Cell Culture in Alginate Microfibers as a Novel Bio-Mimicking Model System for Preclinical Drug Testing. Brain Sciences. 2021; 11(8):1025.

33. Livak KJ, Schmittgen TD. Analysis of relative gene expression data using real-time quantitative PCR and the 2(-Delta Delta C(T)) Method. Methods. 2001; 25(4):402–408.

34. Ayala-Hernández LE, Gallegos A, Schucht P, et al. Optimal Combinations of Chemotherapy and Radiotherapy in Low-Grade Gliomas: A Mathematical Approach. J. Pers. Med. 2021; 11(10):1036.

35. Jiménez-Sánchez J, Martínez-Rubio Á, Popov A, et al. A mesoscopic simulator to uncover heterogeneity and evolutionary dynamics in tumors. PLoS Comput Biol. 2021; 17(2): e1008266.

36. Pérez-Beteta J, Molina-García D, Ortiz-Alhambra JA, et al. Tumor Surface Regularity at MR Imaging Predicts Survival and Response to Surgery in Patients with Glioblastoma. Radiology. 2018; 288(1):218–225.

37. Bette S, Barz M, Wiestler B, et al. Prognostic Value of Tumor Volume in Glioblastoma Patients: Size Also Matters for Patients with Incomplete Resection [published correction appears in *Ann Surg Oncol*. 2018;25(Suppl 3):989]. Ann Surg Oncol. 2018; 25(2):558–564.

38. He Y, Kaina B. Are There Thresholds in Glioblastoma Cell Death Responses Triggered by Temozolomide?. Int J Mol Sci. 2019; 20(7):1562.

39. Bhat KPL, Balasubramaniyan V, Vaillant B, et al. Mesenchymal differentiation mediated by NF-κB promotes radiation resistance in glioblastoma. Cancer Cell. 2013; 24(3):331–46.

40. Furnari FB, Cloughesy TF, Cavenee WK, Mischel PS. Heterogeneity of epidermal growth factor receptor signalling networks in glioblastoma. Nat. Rev. Cancer. 2015; 15:302–310.

41. Zahonero C, Sanchez-Gomez P. EGFR-dependent mechanisms in glioblastoma: Towards a better therapeutic strategy. Cell Mol Life Sci. 2014; 71:3465–3488.

42. Hegi ME, Diserens AC, Gorlia T, et al. MGMT gene silencing and benefit from temozolomide in glioblastoma. New England Journal of Medicine. 2015; 352(10):997–1003.

43. Meng X, Zhao Y, Han B. et al. Dual functionalized brain-targeting nanoinhibitors restrain temozolomide-resistant glioma via attenuating EGFR and MET signaling pathways. Nat Commun. 2020; 11(1):1–15.

44. Barciszewska AM, Gurda D, Głodowicz P, Nowak S, Naskręt-Barciszewska MZ. A New Epigenetic Mechanism of Temozolomide Action in Glioma Cells. PLOS ONE. 2015; 10(8):e0136669.

45. Malta TM, de Souza CF, Sabedot TS, et al. Glioma CpG island methylator phenotype (G-CIMP): biological and clinical implications. Neuro Oncol. 2018; 20(5):608–620.

46. Vaios EJ, Nahed BV, Muzikansky A, Fathi AT, Dietrich J. Bone marrow response as a potential biomarker of outcomes in glioblastoma patients, Journal of Neurosurgery JNS. 2017; 127(1):132–138.

47. Arora A, Somasundaram K. Glioblastoma vs temozolomide: can the red queen race be won?. Cancer Biol Ther. 2019; 20(8):1083–1090.

48. Lee SY. Temozolomide resistance in glioblastoma multiforme. Genes Dis. 2016; 3(3):198–210.

49. Jiapaer S, Furuta T, Tanaka S, Kitabayashi T, Nakada M. Potential Strategies Overcoming the Temozolomide Resistance for Glioblastoma. Neurol Med Chir (Tokyo). 2018; 58(10):405–421.

50. Verhaak RG, Hoadley KA, Purdom E, et al. Integrated genomic analysis identifies clinically relevant subtypes of glioblastoma characterized by abnormalities in PDGFRA, IDH1, EGFR, and NF1. Cancer Cell. 2010;17(1):98–110.

