## Supplementary Material (Text) for "On optimal temozolomide scheduling for slowly growing gliomas"

**Running title:** Optimal temozolomide scheduling for slowly growing gliomas

**Corresponding authors:**

Víctor M. Pérez-García. Mailing address: Edificio Politécnico. Universidad de Castilla-La Mancha. Avenida de Camilo José Cela 3. Phone: +34-926295435. E-Mail:

**Funding:** This research was funded by the James S. Mc. Donnell Foundation (USA) 21st Century Science Initiative in Mathematical and Complex Systems Approaches for Brain Cancer (Collaborative award 220020560, doi:10.37717/220020560); Ministry of Education, Science and Technological Development, Republic of Serbia (ref. number 451-03-9/2021-14/200007); Ministerio de Ciencia e Innovación and FEDER funds, Spain (grant number PID2019-110895RB-I00, doi: 10.13039/501100011033 to VMP-G, and RTI2018-093596 to PS-G); and Universidad de Castilla-La Mancha (grant number 2020-PREDUCLM-15634 to JJ-S).

**Conflict of interests:** The authors declare no competing interests.

**Authorship:** Study design and analysis: PS-G, JJ-S, JS, MP, VMP-G. Writing of manuscript: MP, JJ-S, BS-C, RG, PS-G, JS, VMP-G. Research: BS-C, MD, RG, JJ-S. Mathematical modeling: JJ-S, VMP-G. Software: JJ-S. Murine models: BS-C, RG, PS-G. Cell cultures: MD, MP. Project supervision: PS-G, VMP-G. Funding: PS-G, VMP-G. All authors revised and approved the manuscript.

**Discrete mathematical model**

An adapted version of the on-lattice agent-based mesoscopic model^1^ was used to simulate the glioma longitudinal growth dynamics and its response to the treatment in-silico. The basic cellular agents were clonal populations that could gain or lose cells through the different biological processes incorporated into the mathematical model: mitosis, migration, cell death or trait variations due to phenotypic changes. Cells in the same population behaved in the same way, except for intrinsic noise resulting in stochastic transitions. At each time step, cells could proliferate, die, migrate and/or change their phenotype mimicking gliomas behaviour. Proliferation, death and migration within each population were implemented as in previous works^1,2^. Tumor dynamics were simulated in-silico on a spatial domain discretized on a rectangular grid of voxels (volume units). Dynamics were voxel-specific depending on each voxel and neighbouring voxel occupation. In this work we incorporated three basic cellular populations: proneural cells (either proliferative PNs or quiescent PNq), persister cells (P), and mesenchymal cells (either proliferative MESs or quiescent, MESq). Aiming to reproduce the observed characteristics of both PN and MES phenotypes, proneural cells we assumed to grow faster than mesenchymal cells (25% faster), while mesenchymal cells migrated faster (twice as fast) and were assumed to be less affected by TMZ, to capture their increased resistance.

Since nor in mice models neither in in-vitro experiments there is time for mutational events to play a substantial role in the cellular kinetics, we assumed cells in each compartment to be clonal. PNs and MESs cells were assumed to proliferate, migrate, die or become quiescent with a probability μ_sq_. PNq and MESq cells did neither proliferate nor die, but they were allowed to migrate and revert their phenotypes to the proliferative state with probability μ_qs_. Tumor growth rates and proliferation levels (Ki67) are implicitly determined by the interplay between these two rates.

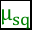

Once a voxel reached a cell number above 70% of its carrying capacity, PNs cells could change their phenotypes to MESs with a rate μ_PT_, due to the effect of local vessel damage and hypoxia. Under exposure to TMZ, PNs cells could either die or enter a persister (P) state with a rate μ_sp_. Cells in P state were allowed to migrate, but neither proliferate nor die. P was assumed to be a reversible intermediate transient state prior to gaining resistance to therapy. If exposure is prolonged, P cells switch their phenotypes to a MESs state with a rate μ_pr_, thus becoming resistant. However, halting exposure to therapy was assumed to allow P cells, but not MES cells, to go back to PNs state at a rate μ_ps_. MESs cells were assumed to be less sensitive to TMZ and could proliferate, migrate, die, or switch to a quiescent state (MESq). MES cells’ transition rates between proliferative and quiescent states are the same as those of PNq cells. Those transitions between the different compartments are summarized in Figure S4.

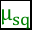

Each spatial voxel may contain several cells belonging to the compartments described before with an upper limit denoted to as local carrying capacity *K*. At a given time step, each cell attempts to perform the different basic processes. These processes can be regarded as two-outcome events, being the possible outcomes success or failure. Therefore, a single cell attempting a given process can be considered a Bernoulli process. Thus, the expected number of successes coming from cells in a given state and a certain voxel attempting a process can be drawn from a binomial distribution with a probability associated to the process. Following this elaboration, the number of cells successfully undergoing division, death, migration or transition to another state are calculated voxel-wise and state-wise at each time step by randomly sampling the corresponding binomial distribution B(N,p), whose *N* will be the number of cells in a given state within a voxel, and whose probability p will be the rate of the process modulated by the time step length Δt.

**Parameter estimation**

To run murine tumor simulations, a grid of 100 x 100 x 100 voxels was used, each voxel having a side length of 100 μm, and a carrying capacity *K* of 200 cells. Therefore, voxel volume was 10^-3^ mm^3^. A time step length of Δt = 15 minutes was chosen, in order to keep a good balance between typical cell dynamics times (proliferation and motility) and to be able to resolve the TMZ exponential decay times of about 2 hours. Simulations started with an initial inoculum in-silico of 3×10^5^ cells, and finished when tumors reached 20 mm^3^, in agreement with typical tumor volumes achieved by the time mice are sacrificed. Therapy was implemented to resemble the actual administration of drug to mice in the experimental part of this study. Three doses of TMZ were simulated in-silico, the first 7 days after initial inoculum. The remaining doses are administered with a gap of 1, 4, 7 or 13 days, depending on the dose spacing selected. TMZ pharmacokinetics were not modelled explicitly; instead, the ratio between current TMZ concentration and TMZ concentration achieving the maximum effect was used. This ratio can range between 0 and 1. TMZ has a plasma half-life of 1.8 hours, which is larger than the selected time step length, so TMZ concentration exponential decay over time can be resolved in the model.

To run human tumor simulations, again a grid of 100 x 100 x 100 voxels was used, but this time voxel side length was increased to 1 mm (with total volume being 1 mm^3^), in order to match typical voxel sizes of high-resolution magnetic resonance images. Carrying capacity was increased to 2 x 10^5^ cells. Time step length was set to Δt = 4 hours, to fasten simulation time while still keeping good resolution of human cell processes. During a 12-hour window starting from each TMZ dose administration, time step length was reduced to 15 minutes, to accurately resolve exponential drug decay. Simulations started with 10 PN cells placed in central voxel. The diagnostic volume was randomly sampled from an empirical distribution of diagnostic sizes obtained from real patient data from TCIA. Unfortunately, an analogous empirical distribution of tumor volumes for sizes at death cannot be obtained, since the last available measurement corresponds to the last imaging follow-up, which does not match patient’s death. To address this issue, we fixed a maximum reachable tumor volume of 120 cm^3^ and simulations were stopped when tumors reached this size. Having defined diagnostic and death volumes in the context of the mathematical model, we could compute a survival time in-silico for every simulated tumor. This provided the required time points in order to run virtual clinical trials.

**Patients**

Volumetric data from a cohort of 69 patients of GBM from The Cancer Imaging Archive (TCIA-TCGA) was used to parametrize the model. Contrast-enhancing volume, necrotic volume and total volume were computed for each tumor in a previous work^3^. With these measures we built an empirical distribution of diagnostic sizes, that we used to random sample realistic diagnostic volumes for simulated tumors. This distribution was validated with independent patient data^4^, where a cohort of 209 GBM patients provided a median preoperative tumor volume of 24.9 cm^3^, and an interquartile range of 11.1-49 cm^3^, which agrees with our empirical distribution.

Mutation, CNV and immunohistochemical data from a cohort of 51 patients of GBM from the TOG study (Therapy Optimization in Glioblastoma) was also used to parametrize the model. TOG study was approved by the Institutional Review Board of all involved hospitals (Marqués de Valdecilla, Sanchinarro, 12 de Octubre, Virgen de la Salud, General Universitario de Ciudad Real, Universitario de Málaga, Manises, Universitario de Albacete). Selection criteria were lack of K27M mutation in H3F3A, lack of V600E mutation in BRAF, IDH1 wild-type tumors, and availability of Ki67 data.

**Virtual clinical trial**

To assess the effect that increasing dose spacing has on survival, we performed a virtual clinical trial with cohorts of simulated human glioblastomas treated under different conditions: control, standard therapy (TMZ during 5 days + 23 resting days), protracted therapies with progressively increasing resting periods, and low dose density therapies with progressively increasing intervals between doses (without resting periods). In standard therapy, 6 cycles were administered, meaning 30 TMZ doses were given. Treatment started 1 week after diagnosis. In all protracted therapies, these 30 doses were spaced either by increasing resting period (but keeping an interval of 1 day between doses), or by increasing interval between doses. We evaluated two types of tumors: fast-growing glioblastomas, with a median Ki67 of 30%, and slow-growing glioblastomas, with a median Ki67 of 10%. 50 tumors were simulated for each branch, so the whole virtual clinical trial comprises 600 simulated tumors. Differences in survival between branches were computed by comparing estimated Kaplan-Meier curves with log-rank test. The median survival differences were also computed for each therapy scheme against control, to assess their effectiveness.

**Parameter analysis of discrete mathematical model**

A two-level parameter exploration was performed with the discrete mathematical model. On the first level, we run virtual murine tumor simulations varying μ_sq_ and μ_qs_. In this way, we generated a cohort of tumors with different Ki67 levels and OS, including fast-growing and slow-growing GBM models. On the second level, we explored the effect of three key parameters: μ_PT_, μ_ps_ and R_fac_. μ_PT_ affects the spontaneous transition from PN to MES cells, thus influencing the time it takes for the PN-MES phenotypic transition to occur within the tumor. μ_ps_ determines the time persister cells can withstand the lack of TMZ before turning back to PNs cells. R_fac_ indicates the ratio between dying fractions of both MES and PN cells when treated when TMZ; hence, a greater R_fac_ means a lower resistance of MES cells compared to PN ones. Considering both levels of the parameter exploration, we generated three cohorts of virtual murine tumors, one per parameter evaluated on the second level. Each tumor was characterized by a triad of μ_sq_, μ_qs_ and μ_PT_/μ_ps_/R_fac_ values. The ranges of values selected for each parameter can be seen at Table 1. Each virtual tumor was simulated five times with the same random number generator seed. In this way, we could isolate the effect of TMZ on survival and MES content, without any other stochastic elements influencing the outcome. More specifically, virtual ‘twinned’ tumors were simulated without treatment (control), and with 1-, 4-, 7- and 13-day spacing.

On the first level of parameter exploration, we reproduced real murine tumor dynamics in terms of OS, Ki67 and PN-MES transition. Parameters μ_sq_ and μ_qs_ were chosen according to a Bayesian criterion. After running several simulations without treatment, those whose initial parameters produced realistic murine tumors (with OS ranging between 30 and 60 days, and Ki67 levels ranging from 5 to 30%) were selected and used to select plausible ranges for these two parameters. This is a rough but effective approach to ABC rejection algorithm. Initial μ_sq_ ranged from 1 to 0.33 days^-1^, while initial μ_qs_ ranged from 0.1 to 0.0166 days^-1^ (Supplementary Figure S5). Tumors with Ki67 levels around 30% have associated an OS of 30 days, thus encompassing the fast-growing group. Meanwhile, tumors with Ki67 around 5% have associated an OS of 60 days, hence belonging to the slow-growing group. Treatment improved survival and reduced MES tumor content, independently of dose spacing.

On the second level of parameter exploration, we observed that increasing the value of μ_PT_ reduces the amount of MES cells in the tumor (in absence of treatment). This effect is more noticeable in fast-growing tumors. When applying TMZ, both 4-day and 13-day spacings showed an increase in MES cells compared to control, being more pronounced for larger values of μ_PT_. However, the increase was smaller for the 13-day spacing (10% more MES cells than control) compared to the 4-day spacing (20% more MES cells than control). Interestingly, some treated slow-growing tumors showed a decrease in MES cells compared to control.

When looking at R_fac_, we observed that, if MES cells are as resistant as PN cells (Rfac = 1), fast-growing tumors are the most benefited from spacings, with the 4-day spacing showing the greatest effect. If MES cells are fully resistant (Rfac = 0), then slow-growing tumors are the most benefited by the spacings, with almost no differences between both 4-day and 13-day spacings. None of these cases are realistic, but they help us narrowing down the behavior it should be expected from a tumor depending on their MES cell content.

Finally, the analysis of μ_ps_ revealed that, for short persistence times, the benefit from spacing is just moderate. However, for long persistence times, OS increase of treated tumors becomes greater, especially for slow-growing ones. Regarding resistance, the greater μ_ps_ is, the larger the increase in MES cells is. 13-day spacing produced a smaller increase in MES cells than 4-day spacing, suggesting that short cycles may be worse when it comes to the emergence of resistance.

**REFERENCES**

1.	Jiménez-Sánchez J, Martínez-Rubio Á, Popov A, et al. A mesoscopic simulator to uncover heterogeneity and evolutionary dynamics in tumors. *PLoS Comput Biol*. 2021; 17(2): e1008266.

2.	Henares-Molina A, Benzekry S, Lara PC, García-Rojo M, Pérez-García VM, Martínez-González A. Non-standard radiotherapy fractionations delay the time to malignant transformation of low-grade gliomas. *PLoS One.* 2017; 12(6):e0178552.

3.	Pérez-Beteta J, Molina-García D, Ortiz-Alhambra JA, et al. Tumor Surface Regularity at MR Imaging Predicts Survival and Response to Surgery in Patients with Glioblastoma. *Radiology*. 2018; 288(1):218-225.

4.	Bette S, Barz M, Wiestler B, et al. Prognostic Value of Tumor Volume in Glioblastoma Patients: Size Also Matters for Patients with Incomplete Resection [published correction appears in *Ann Surg Oncol*. 2018;25(Suppl 3):989]. *Ann Surg Oncol*. 2018; 25(2):558-564.
